# Assessing Codon Language Models for Context-Aware Codon Optimization in Nucleic Acid-Based Medicines

**DOI:** 10.64898/2026.08.11.744178

**Authors:** Shushan Toneyan, Kerstin Scholz, Carlo De Donno, Florian Noack, Simon Auslaender, Tony Cijsouw, Joshua L. Payne

## Abstract

Codon optimization uses synonymous sequence changes to improve the expression and therapeutic performance of nucleic acid-based medicines. Masked language models (MLMs) have recently been proposed as alternatives to traditional, frequency-based codon optimization approaches, yet whether they offer a meaningful advantage over such simpler methods remains unclear. Here we benchmark three prominent MLMs – CaLM, EnCodon and CodonTransformer – across backtranslation fidelity, sequence generation and nine molecular phenotype prediction tasks, and experimentally evaluate model-designed sequences using a secreted embryonic alkaline phosphatase (SEAP) reporter. The models differed markedly in amino-acid fidelity and generated distinct synonymous sequence variants. However, no single model performed best across all benchmark tasks and simple sequence features remained competitive in several settings. Our interpretability analysis revealed that the models integrate a large window of codon context for making predictions, as opposed to frequency-based approaches. Our *in vitro* data showed that MLM-designed variants outperformed conventional and commercial-vendor-derived sequences in both transient and stably integrated expression, supporting the models’ ability to capture translational context beyond codon frequency. Together, our results establish MLMs as effective and complementary tools for codon optimization and suggest that sampling across multiple models may improve the likelihood of identifying high-performing therapeutic sequences.

## Introduction

Nucleic acid-based medicines^1^ are a promising modality for treating and preventing diseases arising from genetic mutations and viral pathogens. This often involves administering a protein-encoding genetic payload, ranging from recombinant DNA delivered by rAAV vectors for sustained protein expression in gene replacement therapies to synthetic mRNA packaged in lipid nanoparticles for rapid immune activation in vaccines. The therapeutic or prophylactic efficacy of these medicines depends heavily on the host cells’ ability to efficiently translate the genetic payload, necessitating precise control over expression levels and translational kinetics. Achieving this control requires the strategic refinement of the genetic payload, particularly through codon optimization, in which synonymous sequence changes are introduced to enhance translational efficiency. By increasing the protein yield per transcript, target expression can be attained at lower administration levels, thereby mitigating the risk of dose-dependent immune responses — such as those triggered by high viral loads or lipid nanoparticle concentrations — and enhancing the overall safety profile of the treatment or vaccine.

Central to the challenge of codon optimization is the astronomical number of possible synonymous DNA or mRNA encodings of any given protein — a combinatorial expansion arising from the degeneracy of the standard genetic code, wherein most amino acids are specified by multiple codons. Despite the equivalence of the final amino acid sequence, these synonymous encodings are not biologically identical, but rather vary in therapeutically relevant phenotypes, most importantly protein abundance and immunogenicity. Variation in protein abundance can arise from the influence of codon choice on mRNA secondary structure, with different synonymous encodings yielding mRNA molecules that vary in stabilities and affinities for post-transcriptional regulators, such as microRNAs^2^ and RNA binding proteins^3^. They can also vary in their translational initiation and elongation rates^4,5^, which can trigger mRNA decay pathways^6^ and influence nascent protein folding^7^. Variation in immunogenicity^8^ can arise because of the presence of particular sequence motifs, such as CpG dinucleotides, which in the context of the rAAV genome can lead to the recognition of pathogen-associated molecular patterns (specifically, PAMP CpG) by innate immune receptors (TLR9)^9.10,11,12^ and ultimately lead to clearing of transduced cells. Synonymous sequence changes can therefore be leveraged to increase the abundance and to reduce the immunogenicity of the payload protein.

Early research on codon optimization relied on heuristics that take into account tRNA gene abundance or the codon frequencies of highly expressed genes, as exemplified by the Codon Adaptation Index (CAI)^13^. While effective at enhancing translation in some contexts and widely used in commercial tools, such frequency-based approaches ignore the important influence of sequence context on the multifaceted determinants of translational efficiency. For instance, maximizing CAI in isolation can inadvertently destabilize mRNA secondary structure, leading to accelerated transcript decay and diminished protein output^14,15^. To address these trade-offs, computational strategies were developed that utilize linear programming to balance codon usage with biophysical constraints, specifically the minimum free energy of the mRNA secondary structure. More recently, supervised deep learning architectures emerged to map coding sequences directly to proxies of translational efficiency, such as ribosomal load^16,17^. However, these approaches either rely on engineered features that fail to capture sequence context, such as CAI, or are constrained by the scarcity of high-quality labeled datasets, especially for synonymous variants, limiting their generalizability across diverse therapeutic contexts.

To overcome the limitations of manual feature engineering and data scarcity, there is a growing shift toward self-supervised learning, which treats genetic sequences as a structured language with their own internal grammar. Masked Language Models (MLMs), in particular, offer a powerful framework for capturing this syntax by training on vast repositories of unlabeled genomic data. By forcing the model to reconstruct masked codons from their surrounding context, MLMs learn to internalize the complex, non-linear dependencies—ranging from local motif preferences to global structural constraints—that define natural coding sequences.

MLMs have been highly successful in various fields, most prominently in natural language modeling^18^ and protein sequence modeling^19^. However, it is unclear how codon MLMs perform in the context of codon optimization. In this study, we benchmark three prominent MLMs for codon optimization: CaLM^20^, Encodon^21^, and CodonTransformer^22^. We evaluate these models on their ability to recapitulate the statistical properties of natural sequences, predict therapeutically relevant phenotypes, and generate diverse sequence libraries for Design-Build-Test-Learn (DBTL) cycles. Using a secreted embryonic alkaline phosphatase (SEAP) reporter, we experimentally demonstrate that MLM-designed sequences outperform frequency-based and commercial vendor-derived sequences in both transient and stably integrated expression. Moreover, we perform interpretability analyses to determine how these models move beyond simple heuristics, revealing that they incorporate local and distal sequence context to propose sequence variants. Our results establish a framework for integrating these models into the rational design of nucleic acid-based medicines, enabling the exploration of a broader and more biologically viable sequence space.

## Results: Model benchmarking

### Models

Existing machine learning approaches to codon optimization include supervised models^23^, causal (autoregressive) models^23,24^, and MLMs^25–27^. Here, we focus on MLMs, which are representation models that can learn codon distributions from unlabeled endogenous sequences and leverage bidirectional sequence context, in contrast to supervised and autoregressive models. These models overcome the issue of data scarcity by leveraging data from endogenous genes of diverse species. We evaluate three prominent MLMs representing different methodological approaches to codon-level representations: CaLM, EnCodon, and CodonTransformer (Table 1).

**Table 1:** Overview of codon MLM architectures and training characteristics.

| Feature | <u>CaLM</u> | <u>cdsFM</u><br>- EnCodon | <u>CodonTransformer</u> |
| --- | --- | --- | --- |
| Tokenization | codon (e.g., ATG) | codon (e.g., ATG) | codon + amino acid (e.g., M_ATG) |
| Species information | Inferred from data (i.e., no species tag) | Inferred from data (i.e., no species tag) | Used as model input during training and sequence generation |
| Fine-tuning | None | Fine-tuned on eukaryotic data | Fine-tuned on genes with top 10% codon similarity index |
| Model size | ~100 million parameters | 90 million , 620 million and 1 billion parameter versions | 89.6 million parameters |
| Input size (tokens) | 1,024 | 2,048 | 2,048 |
| Architecture | Transformer model inspired by ESM | Transformer inspired by RoFormer | Transformer inspired by BigBird |
| Dataset | 9 million cDNA from whole-genome sequencing (European Nucleotide Archive) | 60 million sequences from 5,000 species (NCBI) | 1 million paired coding and amino acid sequences from 164 organisms (NCBI) |
| Sampling | Amino acid not fixed | Amino acid not fixed | Amino acid fixed through tokenization |

CaLM (Codon Adaptation Language Model) was developed as a codon-level alternative to protein language models like ESM and uses codons as tokens. Benchmarks indicate that this representation allows the model to consistently outperform similarly scaled protein language models in predicting phenotypes such as the abundance, subcellular localization, and melting temperature of proteins.

EnCodon, a component of the cdsFM suite, contains models of different scales. In this work we have used the 620 million parameter version as an example of a larger-scale model, which underwent a second adaptation stage on eukaryotic sequences to better capture the nuances of higher-organism codon usage. Like CaLM, EnCodon tokenizes at the codon level without fixing the amino acid identity, requiring additional sampling logic to maintain a target protein sequence during codon optimization.

CodonTransformer, in contrast, utilizes a unique tokenization scheme that represents amino acid–codon pairs as single tokens (e.g., `S_AGC`). This design allows the model to mask only the codon while keeping the amino acid identity fixed (e.g., S_UNK), enabling direct, context-aware back-translation from a protein sequence to a DNA sequence. Unlike the other models, CodonTransformer uses taxonomy IDs as explicit model inputs, allowing for host-specific sequence generation.

How the models specify input and output influences their utility for codon optimization. Although CodonTransformer predicts over the full vocabulary, in practice its amino acid–codon tokenization leads it to assign negligible probability to non-synonymous codons and concentrate probability on synonymous codons. In contrast, CaLM and EnCodon do not encode amino acid identity in the same way and therefore require constrained sampling to generate synonymous variants. This motivates a direct comparison of backtranslation fidelity for CaLM and EnCodon.

### Backtranslation fidelity

We quantified the backtranslation fidelity of CaLM and EnCodon to the reference amino acid sequence by measuring the extent to which the models favor synonymous over non-synonymous substitutions. Such fidelity is important, because the probability mass assigned to non-synonymous codons is not useful for codon optimization. Using 2,012 proteins spanning diverse protein families (Methods), we masked each codon position in turn and computed the normalized probability distribution over the codon vocabulary. Fig. 1a shows two contrasting CaLM outputs, chosen to illustrate model preference for synonymous (Fig. 1a, top) vs. non-synonymous substitutions (Fig. 1a, bottom) of a particular amino acid (Gly, G) at two distinct positions in an example protein; one position shows near perfect backtranslation fidelity where other amino acids’ codons have near zero probabilities, the other displays low backtranslation fidelity where probabilities are spread out across codons encoding different amino acids.

**Figure 1:**
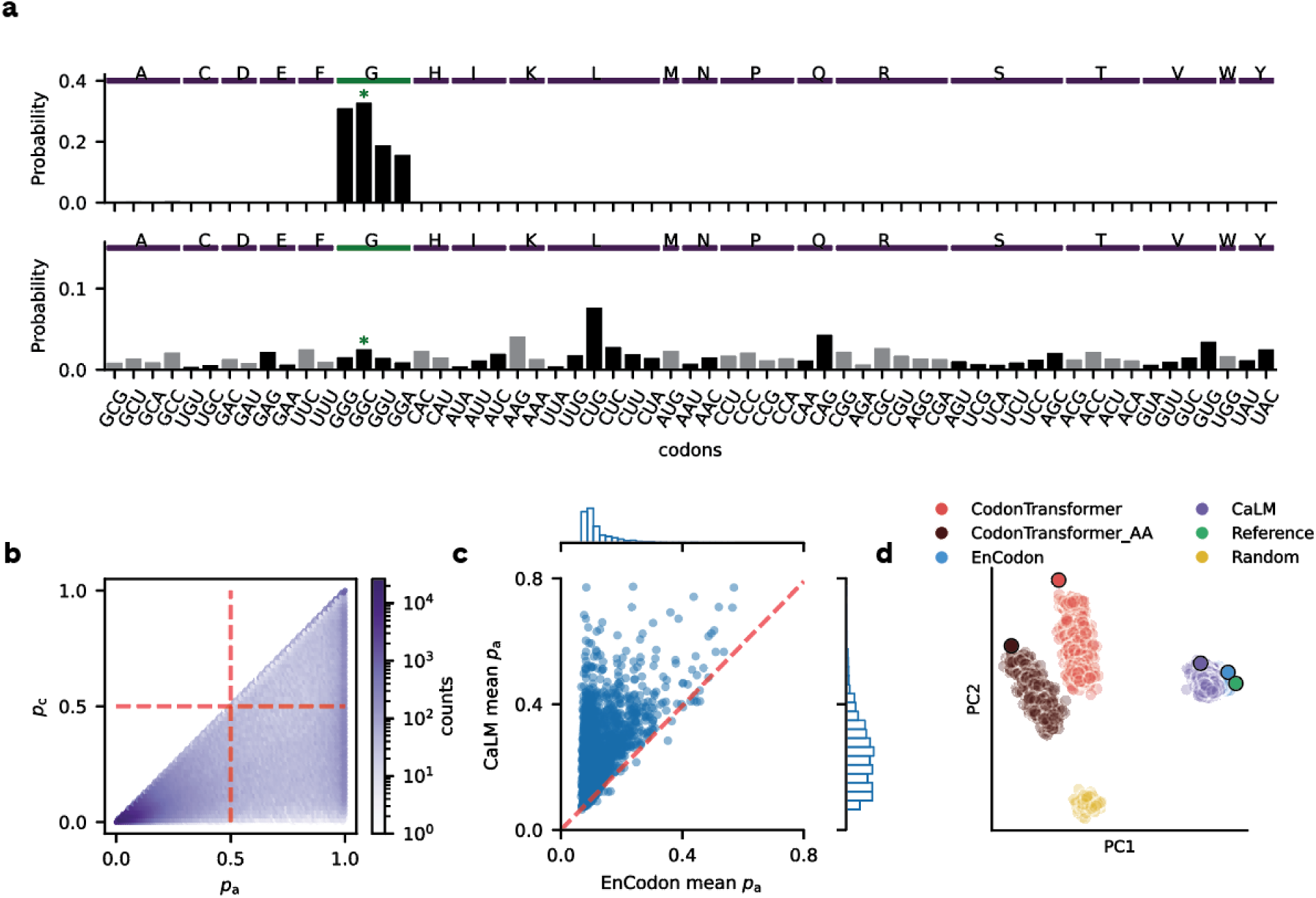
Model evaluation. a, CaLM token probability outputs for the codon GGC at two different positions of the same coding sequence, showcasing distributions with low (top) and high (bottom) heterogeneity. The green bar indicates the reference amino acid and the green asterisk the reference codon. b, Density plot of *p_c_* vs *p_a_* per codon in each of 2,012 proteins using CaLM (n=1,087,855 masked positions). c, Comparison of *p_a_* values between CaLM and EnCodon. Each data point represents the mean *p_a_* averaged across a sequence. d, Variants of a representative human protein, SEAP, including sequences from the three models, random synonymous encodings of the reference sequence, and the reference sequence itself. Each circle depicts a sequence, represented by its nucleotide k-mer frequency (k = 6) and visualized using PCA (n=1,705 points). Darker circles indicate the reference sequence, and the maximum-probability variants from each model. Note the sequence variants from EnCodon and CaLM are overlapping.

To systematically characterize model behavior in the context of backtranslation fidelity across many sequences, we defined two complementary quantities: (i) *p*_c_ - reference codon probability and (ii) *p*_a_ - amino acid fidelity, computed as the sum of the probabilities assigned to the synonymous codons of the reference amino acid. Together, these quantities can be used to identify positions an MLM deems amenable to synonymous change, specifically those with low *p*_c_ and high *p*_a_. This is consistent with the intuition that the model “expects” a different synonymous codon, but not a different amino acid in that sequence context. Fig. 1b and Supplementary Fig. 1 show the density plots of *p*_c_ and *p*_a_ per codon position, across the 2,012 proteins, with the bottom right quadrant representing the positions amenable to codon edits, defined by an arbitrary threshold (*p*_c_<0.5 and *p*_a_ >0.5; see Supplementary Figures 2 and 3 for EnCodon). We observed that the fraction of such positions varies substantially by amino acid, ranging from 2.8% (Glutamine) to 25.2% (Glycine) (Supplementary Fig. 4). This variation remains when controlling for the number of synonyms encoding an amino acid (Supplementary Fig. 4). This shows that (i) *p*_a_ differs across amino acids and sequence contexts, and (ii) the scope for synonymous replacement suggested by MLMs is amino acid–dependent.

To directly compare synonymous substitution preference between CaLM and EnCodon, we repeated the same characterization with EnCodon using the same protein set. CaLM exhibited higher amino acid fidelity than EnCodon (Fig. 1c), with a higher average *p*_a_ per protein in 97% of proteins (1,954 out of 2,012). The mean *p*_a_ across all proteins was 0.246 for CaLM versus 0.124 for EnCodon. These results indicate that CaLM and EnCodon differ substantially in their backtranslation fidelity, highlighting that synonymous substitution preference is an important axis of variation among codon MLMs when used for codon optimization.

### Characterization of edited sequences

Codon MLMs can be used for codon optimization by proposing synonymous edits to coding sequences. Notably, these models have learned the distribution of endogenous coding sequences through self-supervised training. Under the assumption that biologically plausible synonymous codon combinations for a given protein are non-random^28,29^, codon MLMs may help keep edited sequences on the manifold of natural coding sequences. However, because these models are all trained on endogenous genes, it is unclear to what extent different codon MLMs can generate (dis)similar sequences. We therefore set out to analyze edited sequences from each model to test the null hypothesis that these models generate similar, natural-like sequences. This analysis therefore asks whether different codon MLMs explore the same regions of synonymous sequence space or systematically different ones.

To edit sequences using CaLM or EnCodon, we considered each codon position once and in the original sequence context, masking the codon and selecting a potential replacement from the model’s output probability distribution (e.g., Fig. 1a). We only accepted edits that mapped to the reference amino acid and ignored codon positions where non-synonymous edits would be introduced (Methods). An alternative strategy is to set the non-synonymous codon probabilities to zero and sample only from the synonymous codons. Whereas this would increase sequence diversity, it has the drawback of sampling from noisy, low-probability regions of the probability distribution, which often results in the random choice of a synonymous codon. As previously noted, CodonTransformer utilizes a special vocabulary where each token is composed of a combination of amino acid and a codon, e.g. M_AUG or amino acid and an unknown codon e.g. M_UNK. Therefore, the model can edit a position given either only the amino acid context or the amino acid and codon context of the sequence. Using these two strategies we: (i) provide only the amino-acid sequence and infill all codons jointly, or (ii) edit codons one position at a time while providing reference codons at all other positions. When we provide only the amino acid sequence as input, we refer to this setting as CodonTransformer_AA.

To further probe the diversity of generated sequences, we varied the temperature parameter of sampling, i.e., we divided model logits by temperature. Higher temperature settings allow more frequent exploration of codon combinations different from the maximum probability codons generated using lower temperatures. This can be valuable for generating sample diversity for library construction, although the resulting sequences may deviate from the manifold of natural sequences. Therefore, the temperature parameter provides a way to examine the trade-off between sequence diversity and naturalness.

To compare the models in terms of sample generation, we selected 6 proteins (TSP50, CYP24A1, TRIM9, SLC25A3, ADMR, and our wet-lab reporter for benchmarking SEAP) from diverse protein families and edited their coding sequences using CaLM, EnCodon, CodonTransformer and CodonTransformer_AA (Methods). We used four temperature settings ranging from 0.4 to 1 and generated 100 samples per setting. We also included a setting in which the maximum probability codon in each position was selected adding one deterministic sequence variant per model. Additionally, we sampled 100 sequences using random sampling from the pool of synonymous sequences.

To compare the samples of edited sequences, we embedded DNA 6-mer frequencies using PCA (Fig. 1d for SEAP). With the exception of the protein SLC25A3, CaLM and EnCodon sequences occupied partially overlapping regions of the embedding space (Supplementary Fig. 5). The remaining samples formed distinct clusters by model, which differed from the random sample cluster, indicating that different models systematically propose different synonymous substitutions across proteins. Moreover, the relative placement of these clusters in principal component space varied across proteins, highlighting the models’ sensitivity to sequence context in suggesting edits. This is further evidenced by their Hamming distance from the reference sequence, as well as their variation in GC and CpG contents and CAI values (Supplementary Figs. 6-9). The Hamming distance from reference was higher for CodonTransformer compared to CaLM and EnCodon (see Supplementary Fig. 6), which is expected given that CodonTransformer almost never proposes a non-synonymous edit due to its tokenization scheme, so proposed edits are almost always accepted under our sampling scheme. Overall, CaLM samples were slightly further from the reference compared to EnCodon, but the extent of this difference was protein-dependent. We computed the average pairwise codon percentage differences within samples edited with the same model and between different models and found that between-model differences generally exceeded within-model differences (Supplementary Fig 10). Together, these results indicate that different codon MLMs explore distinct regions of synonymous sequence space.

Across models, we observed a drop in GC content with increasing temperature, with most of the samples having GC content above or near the GC content of the reference sequence (Figure ?). Interestingly, different proteins result in different patterns of GC content across the models, providing additional evidence that sequence context plays an important role in codon prediction by the models. Random samples had much lower GC content compared to model-derived samples. Thus, model-based sampling is protein sequence-dependent and leads to sequence properties that differ systematically from those of random synonymous variants. In sum, CaLM, EnCodon, and CodonTransormer generate diverse synonymous sequence variants that vary in their statistical properties, including GC content and distance from the reference sequence, and that are qualitatively different from random synonymous sequence variants.

### Zero-shot prediction of molecular phenotypes

We have shown that these models generate sequence populations that differ from random synonymous sequences and from one another. Next, we characterized whether the learned embeddings of the models aligned with phenotypes relevant to gene therapy and mRNA vaccine design, even though they are not explicitly trained to predict any specific phenotype (e.g., protein abundance, mRNA stability, codon optimality). We assembled a benchmark panel of nine tasks, spanning protein and transcript abundance, mRNA stability, and translational efficiency (Methods; Fig. 2). We note that half of our benchmark pertains to synonymous variants of individual proteins (Fig. 2, shaded region), and all except one task (i.e. mRFP abundance) are from human samples or cell lines. We evaluated representations via linear probing, in which we embedded sequences from a dataset, used PCA to perform dimensionality reduction of features and trained an elastic net model using these features as input to predict the molecular phenotype. These steps were largely replicated from the CaLM study^20^. We used 5-fold cross validation and reported Pearson correlation coefficient (PCC) and compared the results to k-mer frequency baselines computed from nucleotide 6-mers, codon 2-mers and amino-acid 2-mers. In order to test if there is complementary information in different model embeddings, we tested two concatenation schemes. In the first, we concatenated all model embeddings. In the second, we concatenated all model and k-mer embeddings. We then used PCA to reduce the dimensionality ensuring that the number of features in the linear regression model is the same between all comparison points. Together, this benchmark allowed us to test not only the predictive power of individual codon MLMs, but also whether their representations provide information beyond simpler sequence-based baselines.

**Figure 2:**
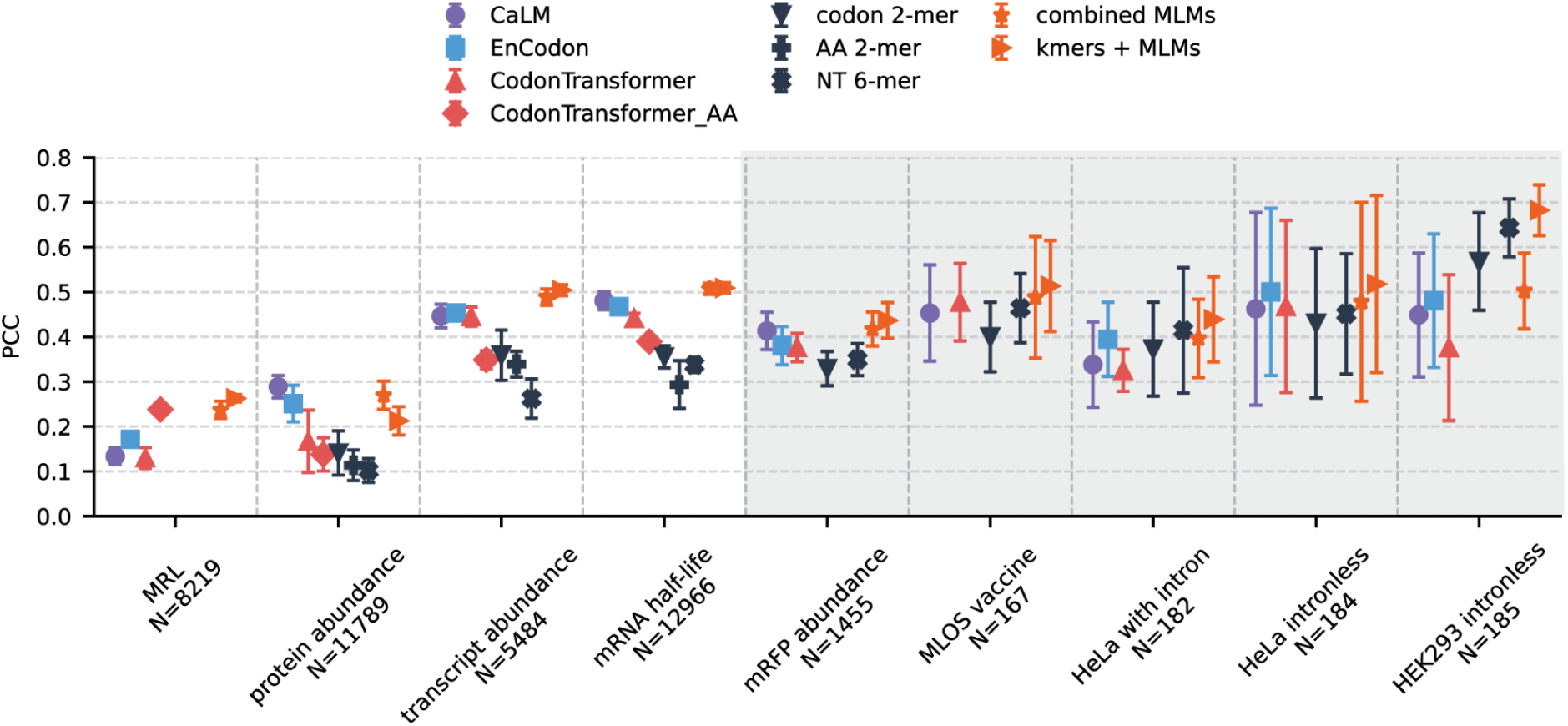
Pearson correlation coefficient for 5-fold cross validation results across the models and baselines for the 9 benchmarking datasets. Symbols represent the mean and error bars the standard deviation across folds. The gray shaded area indicates datasets that include only synonymous variants.

Across most of the tasks, MLM embeddings outperformed the simpler k-mer baselines. The exception is the set of data from Mordenstein *et al*.,^30^ where the simpler k-mer models outperformed all of the MLMs. As the authors note, this could be explained by the GC content of the sequences being correlated with the protein abundance readouts. CaLM and EnCodon performed similarly on most tasks and both outperformed CodonTransformer, except on the mean ribosomal load task where CodonTransformer performed the best. With the exception of the MRL task, Codontransformer performed better given the codon inputs across all the tasks, compared to CodonTransformer_AA. This could be due to the information contained in codons in addition to the amino acid chain that the model has learned or it could be a result of masking every codon leading to the inputs being out of distribution for the model. In all tasks, combining the model embeddings performed as well as or better than the best individual MLM. This implies that the models learn slightly different features and this can be leveraged to get consistently good predictions. Similarly, in all tasks except protein abundance, combining k-mer and MLM embeddings outperformed all other approaches. This shows that MLMs provide additional information on top of the k-mer embeddings and do not simply count k-mers. These results suggest that codon MLM embeddings capture biologically relevant variation and that different models provide complementary predictive signals.

Finally, we analyzed a dataset^31^ of synonymous variants encoding the protein Nanoluc luciferase together with measured protein abundance (Supplementary Fig. 11). For each variant, we computed a pseudo log-likelihood (PLL) score as the negative sum of log probabilities assigned to the observed codons (i.e., an estimate of sequence likelihood under the model). All three models showed predictive signal in this setting, with Pearson correlations (PCCs) of −0.483 (EnCodon; p = 0.005 in Fig. 1f), −0.463 (CodonTransformer), -0.521 (CodonTransformer_AA) and −0.455 (CaLM). Notably, simple sequence-level features were also predictive in this dataset: GC content and codon adaptation index (CAI) yielded PCCs of 0.597 and 0.423, respectively. Additionally, we reanalyzed the datasets from the benchmark panel (grey-shaded tasks in Fig. 2) using PLL directly as a predictor. In these tasks, MLMs performed on par with GC content, suggesting that model-assigned sequence likelihoods capture signals relevant to synonymous sequence function even without supervised fine-tuning. Together, these findings indicate that unsupervised codon MLM scores encode meaningful biological information, although in some settings their predictive power overlaps substantially with simple sequence-level features.

## Interpretability

A key motivation for codon MLMs is that, unlike frequency-based or k-mer approaches, transformer architectures can integrate information across entire sequences. This additional capacity comes at substantial computational and engineering cost, so it is important to understand whether codon MLM predictions actually depend on sequence context in a way that cannot be reduced to local k-mer statistics. In the context of codon optimization, we are interested in two main questions: (1) What determines if the model has a preference towards one or more of an amino acid’s synonymous codons? (2) Why is a model allocating a certain probability mass to a given amino acid? Both of these questions are important for understanding the effect of various sampling strategies and how editing codons in one part of the sequence can influence the model outputs in another part of the sequence^32^.

To answer the first question, we defined a summary statistic called codon preference which describes the heterogeneity in the probability distribution across synonymous codons at a masked position (Methods). This measure takes on a minimum value of 0 when the model assigns equal probability to all synonymous codons and a maximum value of 1 when the model assigns a probability of 1 to a single synonymous codon. We apply this summary statistic to our reference sequences, as well as to perturbed versions of these sequences, in which we corrupt the local or distal sequence context with mask tokens, synonymous codons, or non-synonymous codons. By characterizing how such perturbations change codon preference at masked positions, we gain insight into the sequence determinants of codon choice learned by the models.

### Codon preference in unperturbed sequences

Using the 2,012 proteins, we selected codon positions to mask that adhered to several selection criteria (Methods). Most importantly, we selected codon positions where amino acid fidelity was high (p_a_ >0.8) - resulting in 12% and 2% of positions being selected for CaLM and EnCodon, respectively (Supplementary Fig. 12). We further downsampled these to 14,000 codon positions, 700 per amino acid. Here we focus on results from CaLM, which are qualitatively similar to those observed with EnCodon and CodonTransformer (Supplementary Figs. 13 and 14).

First, we characterized codon preference across the selected positions without any perturbation (Fig. 3a, Supplementary Fig. 15). The distribution of codon preference illustrates that it is more common for the model to have low rather than high codon preference. Specifically, 22% of the positions had codon preference below 0.05, with most of those deriving from amino acids encoded by 2 synonymous codons (69%) (Supplementary Fig. 15). Such positions where the model displays little to no preference for particular codons may be amenable to synonymous codon change for the purpose of increasing sequence diversity in experimental libraries or depleting unwanted sequence motifs, such as CpG dinucleotides. In contrast, we reason that at positions with higher codon preference, the model is more likely to suggest a synonymous change that is functionally relevant. Such positions are in the minority. For example, only 35% of positions have a codon preference higher than 0.25. Moreover, as codon preference increases, so does the probability that the model’s top recommendation is the reference codon rather than a synonym (Fig. 3a inset). Indeed, the model recommends a synonym to the reference codon in only 20% of positions with codon preference above 0.25. Together, these results suggest that codon optimization using CaLM or EnCodon will result in only a small number of sequence edits relative to the reference sequence, because positions favorable to synonymous sequence changes (i.e., high amino acid fidelity, high codon preference, and high probability for a synonym of the reference codon) are concentrated at a relatively small subset of codon positions. We next asked what features of sequence context shape these model preferences.

**Figure 3:**
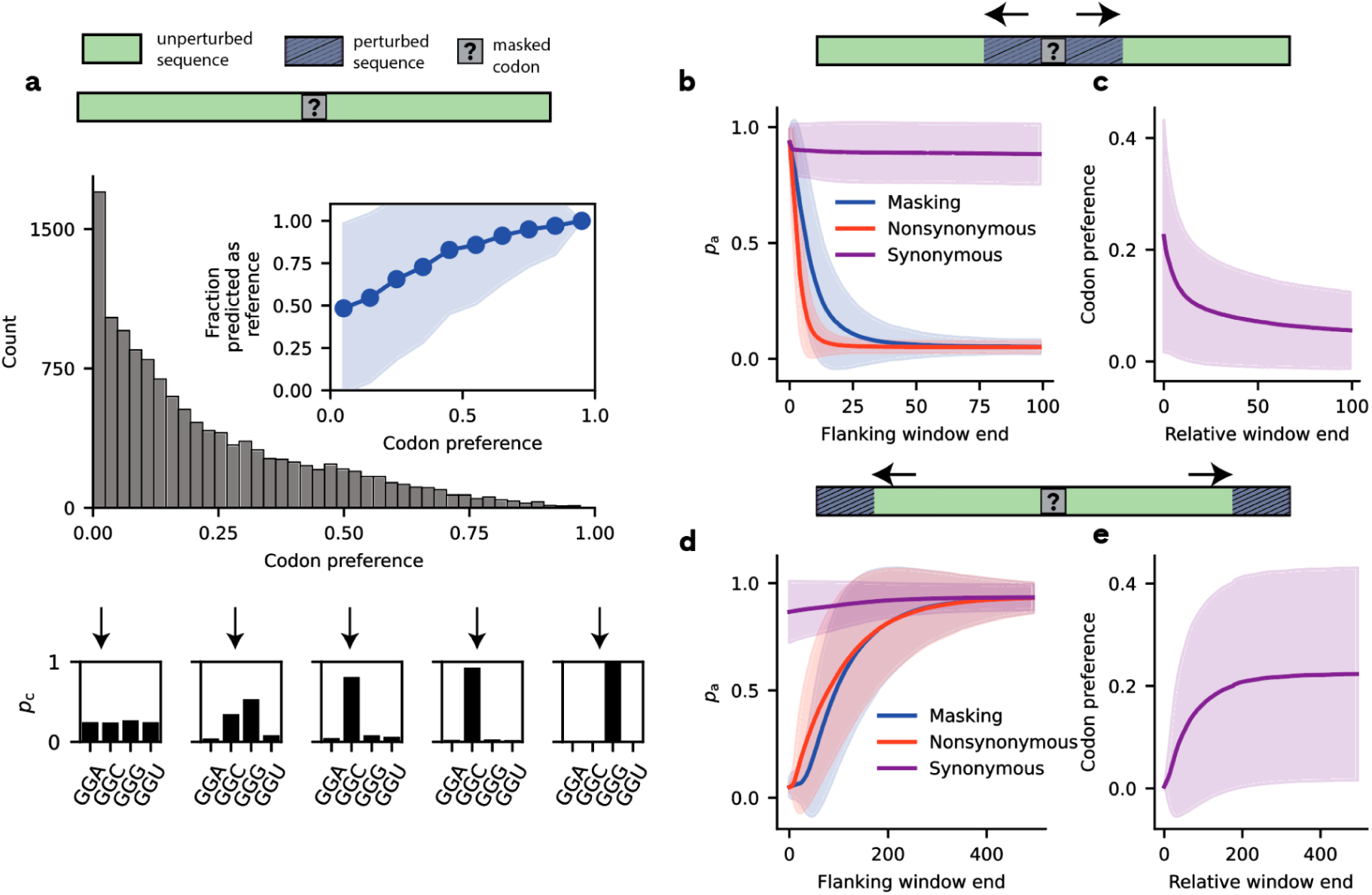
Model sensitivity to context perturbations. a, Codon preference distribution in unperturbed sequences. The top diagram shows a schematic representation of the analysis - i.e., recording codon preferences in reference sequences by masking a codon position. The histogram shows the distribution of codon preference values for 12,600 codon positions where the input codon has a synonym (i.e., excluding the 1,400 Tryptophan or Methionine codons). The inset shows the mean fraction of positions where the codon with the predicted highest probability corresponds to the input codon. The shaded area shows the standard deviation around the mean. The lower panel shows examples of synonymous codon probabilities (normalized to sum up to 1) resulting in codon preference values close to 0, 0.25, 0.50, 0.75 and 1.0 respectively. b, Local context perturbations. The top panel shows a schematic of the perturbation analysis conducted where the local context around the masked codon is perturbed with one of the 3 perturbation types: masking, non-synonymous substitutions, or synonymous substitutions. The line plot shows amino acid fidelity, p_a_, in relation to the window size of the perturbation, stratified by perturbation type. c, The effect of local synonymous perturbation on codon preference. The line plot shows the change in codon preference with an increasing synonymous perturbation window. d, Distal context perturbation. The top panel shows a schematic of the analysis conducted where the unperturbed local context around the masked codon is flanked symmetrically by a distal perturbed sequence. The line plot shows p_a_ in relation to the window size of the perturbation,stratified by perturbation type. e, The effect of distal synonymous perturbation on codon preference. The line plot shows codon preference in relation to the window size of synonymous perturbation. In panels b-d, solid lines represent the mean, and shaded areas the standard deviation, across codon positions. Panels b and d pertain to data from all 14,000 sequence positions selected such that p_a_ is above 0.8 and c and e to 12,600 sequence positions. Note that for local perturbations (panels b, c), the window end refers to the perturbation area, whereas for distal perturbations (panels d, e), the window end refers to the unperturbed area. All data pertain to CaLM.

### What influences model predictions?

The development of codon MLMs is often motivated by application to codon optimization for nucleic acid based medicines^21,22,25^. However, these models regularly recommend non-synonymous sequence changes (see Fig. 1a). To understand the factors shaping model preference for an amino acid at a given position, we investigated both local and distal context importance. We investigated local context importance by symmetrically perturbing the flanking sequence around selected codon positions using various window sizes. By performing local sequence perturbations, we analyzed if local sequence context is necessary for the model predictions or if it is redundant with the distal context. Distal sequence context importance was characterized by symmetrically corrupting various window sizes, but leaving the sequence immediately flanking the masked codon intact. Through distal sequence perturbations, we only allow the model access to the local sequence context, thus testing if local sequence context is sufficient to recover the predictions. To disentangle the effects of amino acids and codons in a sequence context, we applied three perturbation types with increasing severity of information corruption:

i. *Synonymous perturbations*: preserving amino acid identity to determine whether synonymous codon choices affect predictions.
ii. *Masking perturbations*: providing no information about codon or amino acid identity at perturbed positions.
iii. *Non-synonymous perturbations*: changing the amino acid context to test whether amino acid identity influences model output. In this case, the information about the amino acid neighborhood of the selected codon is incorrect.

### Local context perturbations

To evaluate local context importance, or necessity, we systematically perturbed progressively larger windows (ranging from 1 to 100 codons) flanking a selected, masked codon, using each perturbation method. This evaluation was then repeated for all eligible codon positions in the protein set. Fig. 3b shows that amino acid fidelity, *p*_a_, decreases monotonically with window size for all three perturbation types. For synonymous perturbations, *p*_a_ decreases to a minimum of only 95% of its value when unperturbed, indicating that synonymous codon context locally has a negligible impact on model preference for amino acid identity. In contrast, *p*_a_ decreases rapidly with increasing window size for both masking and non-synonymous perturbations. For example, *p*_a_ dropped below 20% of its value when unperturbed at a window size of 17 using masking perturbation and at a window size of only 7 using non-synonymous perturbation. The more severe drop under non-synonymous perturbation highlights the governing influence of flanking amino acid identity on model predictions. This suggests that model predictions of amino acid identity are primarily influenced by flanking amino acid identity and are nearly agnostic to which synonym encodes those amino acids.

### Distal context perturbations

The previous analysis suggests that local sequence context is necessary for the models to correctly predict the reference amino acid at a masked position, but does distal context also play a role? In other words, is the local context itself sufficient for the model to correctly predict amino acid identity at masked positions, or is distal sequence also required? To address these questions, we ran a complementary set of analyses (Fig. 3d). We explored the sufficiency of the local sequence context for correctly predicting amino acid identity by restoring a progressively longer context of reference sequence around a selected codon in an otherwise perturbed background. In the most extreme case, we unmasked only 1 codon flanking the masked position and asked if the codon identity of these immediate neighbors is sufficient to correctly predict the masked codon. If the model could make accurate predictions with such a minimal context window, it would suggest that the model relies solely on the local neighborhood and does not incorporate information from the broader sequence context. In this scenario, the model’s behavior would be analogous to that of a k-mer model for codons. Conversely, if the local neighborhood is insufficient for predicting the reference amino acid or codon, it would indicate that the model is capturing more complex interactions that extend beyond the immediate vicinity of the masked position.

Starting with a completely perturbed sequence, we gradually recovered the reference sequence in a window around the masked codon with a step size of 5 and a maximum window of 500 on each side flanking the masked codon, recording amino acid fidelity at each step. Similar to our analysis of necessity, synonymous perturbations had almost no effect on *p*_a_, even at the extreme of synonymously recoding every position in the protein. Specifically, *p*_a_ dropped from 0.93 at the maximum window size to only 0.87 when the entire sequence context around the masked position was synonymously recoded. In contrast to our analysis of necessity, masking and non-synonymous perturbation are roughly equivalent in their effect on model preference for the reference amino acid. Under masking and non-synonymous perturbation, *p*_a_ recovered to 50% of its maximum value at local window sizes of 111 and 95, respectively, whereas in our necessity analysis it dropped sharply at smaller window sizes. This reveals that while local sequence context is necessary, it is not sufficient for predicting reference amino acid identity in isolation from the remaining sequence context. Together, these findings indicate that CaLM, EnCodon, and CodonTransformer integrate both local and distal context to predict masked codons, with local sequence providing the dominant signal and distal sequence contributing additional, complementary information (Supplementary Figs. 13 and 14). This distinguishes codon MLMs from simpler k-mer-based approaches, which rely only on the immediate sequence neighborhood.

### Context-dependence of codon preference

From our previous analyses, it is evident that synonymous codon perturbations, local or distal, have only a marginal effect on amino acid fidelity at masked positions (Fig. 3 b and d - purple lines). However, such perturbations may nonetheless affect the synonymous codon probabilities at the masked position. To determine if this is the case, we quantified codon preference, in response to both local and distal synonymous codon perturbations, using increasing window sizes. Fig. 3c shows codon preference decreasing with increasing local window size, with codon preference dropping by 50% relative to no perturbation at a window size of only 13 codons. Synonymous codons immediately surrounding the masked position therefore strongly influence model preference for particular codons at the masked position, in contrast to model preference for the reference amino acid at the masked position. Additionally, Fig. 3e shows that codon preference relies on synonymous codon choice distal to the masked position, with codon preference reaching 50% of its value without perturbation at a window size of 55 codons. These results indicate that the model relies on both the local and distal sequence context and that it is sensitive to which synonyms are used to encode a fixed amino acid chain.

In contrast to autoregressive approaches to sequence design, which generate DNA or mRNA sequences iteratively from 5’ to 3’, codon MLMs use the full length of a coding sequence to make predictions for masked positions. To understand if codon MLMs take advantage of sequence information on both sides of the masked position, we repeated our perturbation analyses but only perturbed the sequence upstream or downstream of the masked position. Supplementary Fig. 16a shows the summary of results for CaLM. In general, the model relies on sequence information on both sides of the masked position, such that, on average, codon preference decreases symmetrically as either the upstream or downstream sequence is perturbed. In contrast to our previous analyses using bi-directional perturbations, a larger window size of ∼70 is needed for codon preference to drop below 50% of its value without perturbation. This indicates that pertinent information is encoded on both sides of the masked position. To determine if model predictions at masked positions sometimes depend more heavily on upstream or downstream sequence, we plotted codon preference at the masked position when perturbing the upstream sequence in relation to codon preference at the same position when perturbing the downstream sequence (Supplementary Fig. 16b and c). In this plot, large deviations from the identity line indicate predictions at masked positions that depend asymmetrically on sequence context. Whereas such deviations are the exception, 18% of positions showed an absolute difference in codon preference greater than 0.1 between upstream and downstream perturbations (Supplementary Fig. 16b). This suggests that access to the full sequence context is practically useful, because codon preferences at a subset of positions cannot be recovered from one-sided local information alone.

## Results: SEAP Screen

We established a sequence design and validation framework to systematically benchmark the ability of MLMs to optimize protein output by codon optimization. This framework utilizes Secreted Embryonic Alkaline Phosphatase (SEAP) — a reporter enzyme derived from human placental alkaline phosphatase (PLAP) and based on a variant previously developed by Kaberniuk and colleagues^33^ — as a sensitive reporter system (Fig. 4a). Because SEAP is actively secreted into the culture supernatant, it permits non-destructive, high-throughput kinetic profiling of translational efficiency, while its intrinsic heat stability ensures a high signal-to-noise ratio and a wide linear dynamic range capable of resolving expression variations induced by codon optimization^34^.

**Figure 4:**
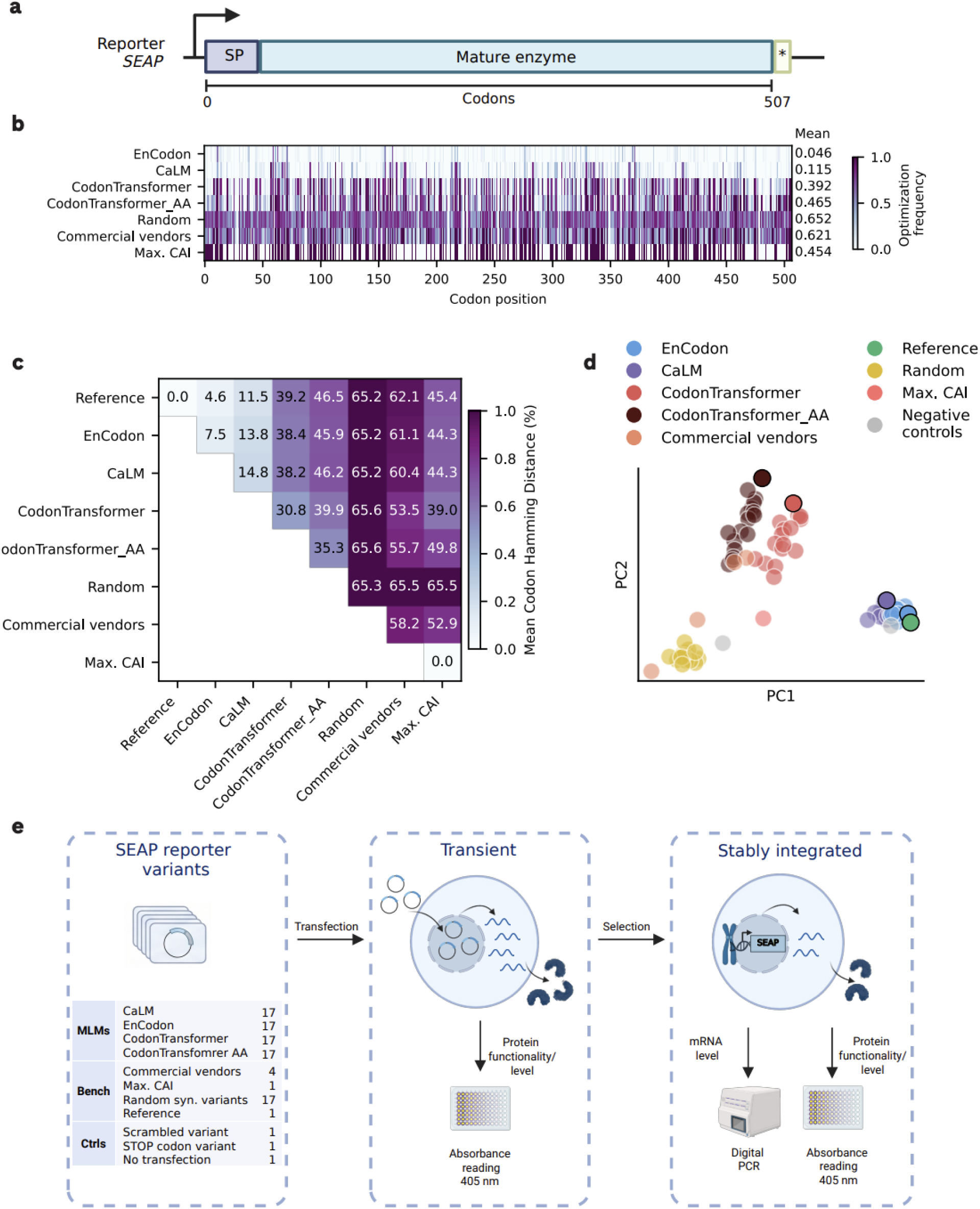
*In silico* design of the human SEAP reporter framework and sequence analysis of generative codon optimization models. a, Schematic representation of the modified human secreted embryonic alkaline phosphatase (SEAP) reporter framework utilized for arrayed *in vitro* benchmarking of MLM-derived codon optimized sequences.

To assess the efficacy of codon optimization by MLMs, we designed a library consisting of 68 sequences generated by the four distinct MLMs previously described. For each model, we generated 17 unique randomly sampled variants across a sampling temperature range of T = 0.0 to 1.0. We constrained sequence design to synonymous codon usage, intentionally omitting secondary filters based on intrinsic sequence features or predictive computational models of molecular phenotypes, thus testing these MLMs “out of the box”.

The expression cassette is driven by a constitutive shortened human elongation factor 1-alpha (EF1alpha) promoter. The encoded reporter is derived from human placental alkaline phosphatase (PLAP), based on a framework established by Kaberniuk *et al.* (2021). To ensure efficient extracellular secretion of the mature enzyme, the reporter features a modified N-terminal signal peptide (SP) optimized via deletion of the first five amino acids, coupled with the complete removal of the native C-terminal glycosylphosphatidylinositol (GPI) anchor signaling sequence. b, Each row represents one SEAP variant grouped by optimization model; each column represents one of 507 codon positions. Purple pixels indicate codons that differ from the reference sequence (top row; 0% edits by definition) and white pixels represent conserved codons. The mean percentage of edited codons per group is shown at right. c, Pairwise mean codon Hamming distance (%) between optimization models. Each cell shows the average fraction of codons that differ between all variant pairs from two groups. Within-group distances reflect sequence diversity among variants generated by the same model across all temperatures. d, Principal Component Analysis (PCA) of the *in silico* variant library based on nucleotide 6-mer frequencies. Synonymous SEAP variants (n=93) project into distinct clusters segregated by generative model type, reflecting model-specific sequence characteristics. Commercial variants display a broader distribution across the embedding space. e, We outline the two-staged experimental screening workflow for *in vitro* validation of codon optimized SEAP sequences. We first transiently tested all MLM derived sequences (n=68; 17 per model) and benchmarks (commercial vendor derived sequences, n=4; max. CAI, n=1; Random synonymous variants, n=17; reference, n=1), as well as controls (scrambled variant, n=1; stop codon variant, n=1; no transfection control, n=1). As a read-out, we determined kinetic SEAP activity in the supernatant (via colorimetric pNPP substrate conversion measured at OD_405_) to assess protein levels. Formally, the colorimetric absorbance assay measures properly secreted, functional SEAP activity, which we interpret as SEAP levels (i.e., assuming that codon usage-dependent misfolding ^35^, which may affect SEAP secretion and/or function, is negligible). Subsequently, we integrated these variants site-specifically as a single copy to mimic low-copy therapeutic delivery. Characterization couples kinetic SEAP measurements with absolute transcript quantification via digital PCR (dPCR) to determine single-transcript specific productivity.

We contrasted the MLM-derived cohort against a baseline of traditional benchmarks and controls (n = 26). This baseline included the Kaberniuk-derived reference (variant SEAP_89), a maximum CAI variant, and four sequences that commercial vendor algorithms (Twist Bioscience, Genewiz, GeneArt GeneOptimizer, and IDT) produced using either DNA sequence optimization or amino acid back-translation pipelines. Further controls consisted of 17 random synonymous codon sequences (Random) and negative controls, including a scrambled sequence, a stop variant, and non-transfected cells (Supplementary Table 1).

To assess how broadly each MLM explores synonymous sequence space, we created a codon edit map (Fig. 4b, Supplementary Fig. 17) displaying across all variants (and temperatures) for each model or method the optimization frequency across the 507 codon positions of SEAP. First, we expected EnCodon and CaLM to produce sequences that remain close to the Reference, because of the sampling approach, which ignores codons positions where non-synonymous edits are proposed; as described above EnCodon and CaLM frequently propose such edits, in contrast to CodonTransformer or CodonTransformer_AA. Indeed, with our sampling strategy EnCodon and CaLM produce mean editing frequencies of 4.6% and 11.5% of the 507 codon positions, respectively, compared to 39.2% for CodonTransformer and 46.5% for CodonTransformer_AA. Next, we computed mean pairwise codon Hamming distances between all model (and method) groups (Fig. 4c). Here, we observe that the four commercial vendors tested generate diverse sequences between each other (within-group Hamming distance of 58.2%), in line with previous findings^24^, and compared with the models (53.5-62.1%). As expected, randomly generated sequences are the most diverse within-group, and compared to the sequences generated by all other models and methods. Interestingly, the sequence with max CAI shows a similarly high Hamming distance from all other methods/models suggesting that these do not optimize for CAI alone. The within-group Hamming distances for Encodon and CaLM (7.5% and 14.8%, respectively) are the lowest across all groups, as is the Hamming distance between Encodon and CaLM (13.8%). Together this shows that CaLM and EnCodon sample from a relatively narrow range of diverse sequence alternatives near to the reference. In contrast, CodonTransformer, and especially CodonTransformer_AA, explore most broadly across sequence space.

To further characterize the library sequences, we mapped all variants via PCA based on 6-mer nucleotide frequencies (Fig. 4d). As observed previously (Fig. 1d), sequences clustered by model type, with each model forming a distinct cluster, whereas commercial variants were more broadly distributed across the embedding space. We also observed that these clusters differed in GC content. This suggests that the library is diverse both in k-mer usage and GC content.

We tested the variant library in a two-stage pipeline (Fig. 4e): first, transient expression in engineered HEK293A cells to measure secreted SEAP levels (absorbance readout) at 24-hours post-transfection, then subsequent single-copy genomic integration to assess long-term expression and eliminate plasmid copy-number effects. Transcript levels generally do not strongly correlate with protein concentration^36^ however can be impacted by codon optimization^37^. Therefore, in the stably integrated context, we quantified both protein and mRNA levels to determine if MLM-based methods can optimize for transcription and/or translation^37^. The ratio of relative protein output and relative RNA abundance yields specific productivity — e.g., low mRNA paired with high protein indicates efficient translation^38^. This *in silico*–to–*in vitro* framework provides a scalable benchmark for evaluating codon optimization strategies.

In the transient readout, MLM variants produced significantly higher relative SEAP activity than Random (median 1.76 vs. −0.31; Tukey HSD, p < 0.001) and Reference (t-test, Bonferroni-corrected, p < 0.001), with 83.8% of variants exceeding the reference, compared to 5.9% of Random variants (Fig. 5a, Supplementary Fig. 18a). EnCodon had the most variants exceeding reference (17/17), followed by CodonTransformer (15/17), CaLM (14/17), and CodonTransformer_AA (11/17).

**Figure 5:**
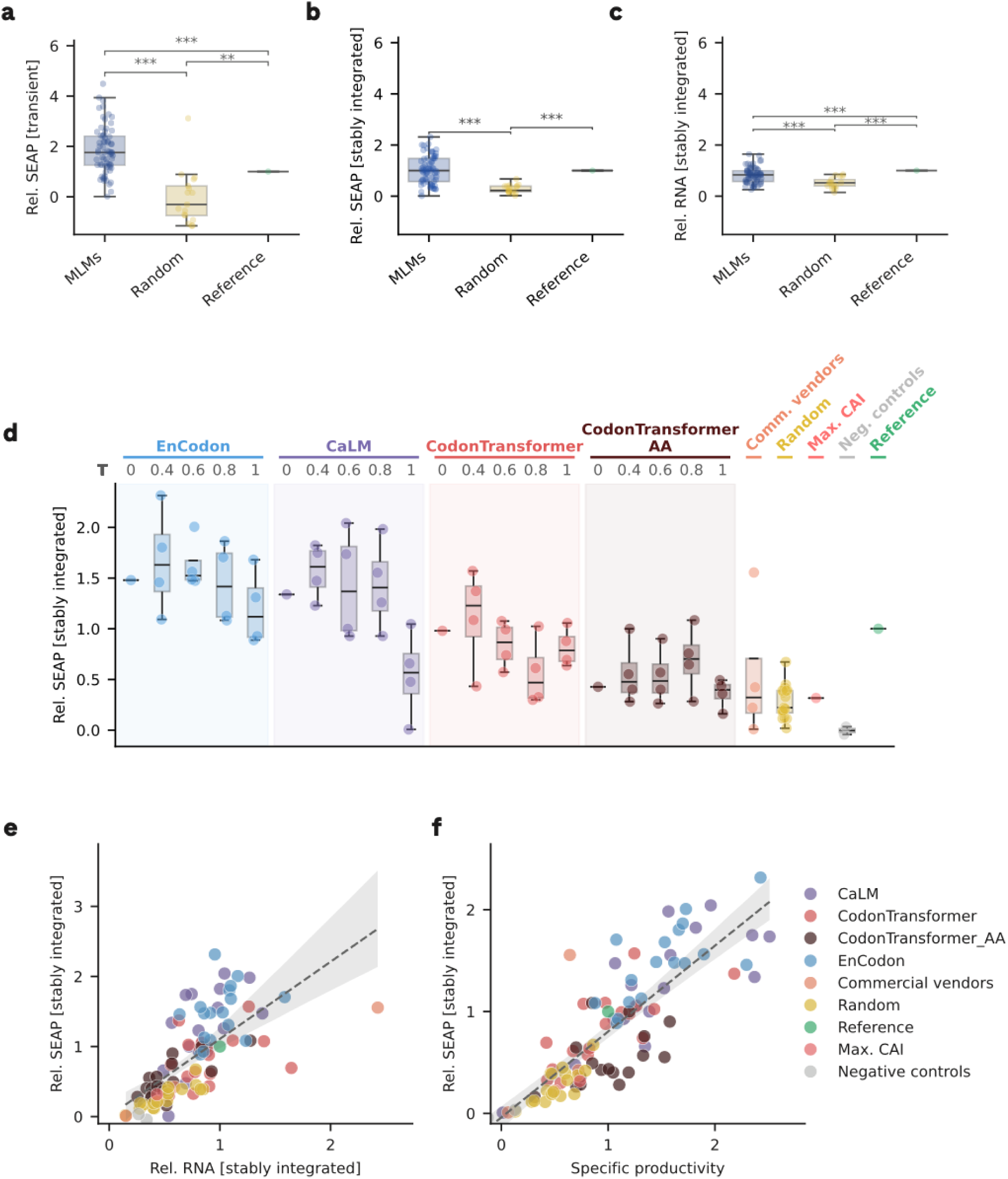
*In vitro* benchmarking and translational evaluation of codon language models across transient and stably integrated platforms. a-c, Global benchmark comparison of relative SEAP activity across MLM-generated sequences (n=17), random synonymous sequences (Random) (n=17), and the reference construct (n=1) in transient (a), stably integrated (b) expression and relative RNA levels (c). Boxplots show median and interquartile range of the group; individual data points represent each variant within the group. Statistical comparisons: Tukey HSD for groups with n ≥ 2; one-sample t-test (Bonferroni-corrected for the number of comparisons involving the reference) for group level-comparisons against reference (n=1) (*p < 0.05, **p < 0.01, ***p < 0.001). d, Stably integrated SEAP expression levels categorized by MLM and sampling temperature (ranging from T = 0.0 to 1.0). Each boxplot illustrates the distribution for a single model–temperature pair, with n = 1–4 variants per condition. Benchmark categories, including Random sequences, Commercial Vendors, Negative Controls, and the Reference construct, are situated to the right. The identity of each model is represented by distinct colored bands. e, Relative RNA expression versus relative stably integrated SEAP activity (n = 93 variants). Dashed line, linear regression with 95% CI. f, Specific productivity versus stably integrated SEAP expression. Higher specific productivity indicates greater translational output per unit mRNA. All data are expressed as the mean ± SD of *n* = 3 independent biological replicates, with the exception of the relative RNA of SEAP_71, where *n* = 2 due to a failed extraction.

After genomic integration this trend persisted, though at lower levels (Fig. 5b, Supplementary Fig. 18b). MLMs outperformed Random (median 1.00 vs. 0.22; Tukey HSD, p = 0.013), but did not differ from Reference, on average (p = 0.73), with only 50.0% of MLM variants exceeding Reference (as compared to 0% of Random variants). EnCodon again led (15/17), followed by CaLM (12/17), CodonTransformer (6/17), and CodonTransformer_AA (1/17) (Fig. 5b). Transient and stably integrated expression correlated positively across the 93 variants (Spearman ρ = 0.741; R² = 0.482; Supplementary Fig. 19).

Stably integrated SEAP expression varied significantly across the four MLMs (ANOVA F = 6.1, p < 0.001; Fig. 5d). To assess whether sampling temperature had an effect on expression within each model, we performed separate one-way ANOVA per model. Only CaLM showed a significant temperature effect (ANOVA F = 5.3, p = 0.015), driven by reduced expression at T=1.0. EnCodon, CodonTransformer, and CodonTransformer_AA showed no temperature dependence (p > 0.05), though the limited number of variants per temperature reduces statistical power to detect subtle effects. In the transient assay, no significant differences were detected across MLM conditions (ANOVA F = 1.6, p = 0.11). Notably, MLM variants significantly outperformed both commercial vendor-optimized sequences (Tukey HSD, p < 0.001) and the max. CAI construct (one-sample t-test, Bonferroni-corrected, p < 0.001) in transient expression. In the stably integrated context, the measured activity of all MLM-derived variants remained significantly higher than max. CAI (p = 1.5 × 10⁻⁵). The comparison with commercial vendors did not reach statistical significance (Tukey HSD, p = 0.41), likely due to the small sample size of commercial vendor variants (n=4), though MLMs showed a higher median expression.

Relative RNA levels correlated positively with stably integrated SEAP output (Spearman ρ = 0.77, R² = 0.44; Fig. 5e), confirming a transcriptional contribution to protein expression differences^39^. However, we observed that specific productivity — defined as the ratio of relative protein output to relative RNA abundance — showed a stronger association with protein expression (Spearman ρ = 0.83, R² = 0.68; Fig. 5f). Additionally, relative RNA level medians of all MLM, random and commercial vendor variants are below the reference (Supplementary Fig. 18c). This indicates that high-performing variants are not only transcribed at higher levels but also translated more efficiently per mRNA molecule, potentially due to improved mRNA stability or enhanced translational initiation or elongation rates^39^. Overall, MLM variants achieved a median specific productivity of 1.20 (max ∼2.5), compared with 0.47 for random and 0.57 for commercial vendor sequences.

## Discussion

Codon optimization remains a central challenge in the development of nucleic acid-based medicines because the same protein can be encoded by an astronomical number of synonymous mRNA sequences, and these sequences can differ in therapeutically relevant molecular phenotypes. Although frequency-based codon optimization methods have been used for several decades, machine learning-based approaches — and, more recently, language models — have emerged as alternative strategies to address this problem. However, the proliferation of such models, particularly masked language models (MLMs), has created a benchmarking gap, leaving it unclear which models offer the greatest utility for the design of nucleic acid-based medicines. Moreover, we lack a mechanistic understanding of how these models integrate local and distal sequence context to differentiate themselves from traditional, frequency-based methods. Here, we fill these knowledge gaps by benchmarking and performing interpretability analyses on three prominent codon MLMs — CaLM, EnCodon, and CodonTransformer — and by experimentally validating their sequence designs in an *in vitro* arrayed expression screen.

Overall, our results establish codon MLMs as promising tools for therapeutic sequence optimization. Linear probing of model embeddings generally revealed increased predictive power relative to frequency-based methods on tasks relevant to transcription, mRNA stability, and translation for both nonsynonymous and synonymous sequence variants. However, in five out of nine tasks, simpler models based on k-mer statistics or other sequence features, such as GC content, were equally predictive — a gap we anticipate could be narrowed through fine-tuning on labelled datasets. We did not observe a single model that consistently outperformed all others across computational benchmarks, and the sequences generated by each model are qualitatively distinct, indicating utility in sampling from multiple models to maximize sequence diversity in library-based experiments. Combining model embeddings and k-mer frequencies yielded the best performing model across datasets. These findings suggest that the models have overlapping but distinct strengths, better understood in a task-dependent framework than through a simple overall ranking^40^.

This interpretation is reinforced by our *in vitro* arrayed screen, in which SEAP served as a reporter for codon-dependent expression differences. MLM-designed sequences consistently outperformed random synonymous codon assignment and, in transient expression, significantly outperformed the reference sequence. Among the four models, EnCodon achieved the most consistent performance, with all 17 variants surpassing the reference in transient expression and 88% in stably integrated expression, followed by CaLM (82% and 71%) and CodonTransformer (88% and 35%). CodonTransformer_AA showed the weakest performance, particularly in stably integrated expression where only 1 of 17 variants exceeded the reference. Commercial vendor-optimized sequences and the sequence designed to maximize CAI tended to have lower expression than the reference sequence, with only a single commercial variant exceeding it. In summary, the variation across models supports a strategy of generating candidates from multiple MLMs and screening empirically, leveraging the complementary sequence space each model explores — reflected in distinct codon Hamming distances, GC content profiles, and MFE distributions.

The experimental data suggest that the benefit of codon optimization is primarily translational: specific productivity explained 68% of stable SEAP expression variance compared with 44% for mRNA levels alone, indicating that high-performing variants are translated more efficiently rather than simply being more abundant at the transcript level. When we correlated key sequence descriptors—including GC3, CAI, CSI, rare codon frequency, MFE, predicted mRNA half-life and pseudo-log-likelihood (PLL) scores from each MLM—with expression, we initially identified GC content as the dominant confounder, but this association was driven by the inclusion of the low-performing random control group, which has both low GC content and poor expression; excluding this group eliminated GC content as the leading predictor, a pattern consistent with Simpson’s paradox, demonstrating that MLM-driven optimization captures coding rules beyond simple GC bias (Supplementary Fig. 20). To further assess potential confounding effects, we used OpenSpliceAI to screen for cryptic splice sites introduced by synonymous substitutions. Predicted splice sites were detected in one random variant, one commercial vendor variant, and the max. CAI construct, which may partially account for their reduced expression (Supplementary Fig. 21). Notably, all MLM-generated sequences were free of predicted splice sites, suggesting that these models implicitly learn to avoid splicing-competent motifs ^39,41–45^.

An effective codon optimization method must satisfy several requirements simultaneously: it should preserve the encoded amino acid sequence, produce sequences with favourable molecular phenotypes, maintain global sequence properties such as host-compatible GC content, avoid undesirable motifs such as CpG dinucleotides when relevant, and ideally generate a diverse pool of candidate sequences rather than a single design. Our results suggest that codon MLMs satisfy some, but not all, of these requirements in their current form. For example, only CodonTransformer’s tokenization allows for seamless backtranslation of sequences. CaLM and EnCodon require additional sampling procedures to enforce synonymous editing, which can affect the outputs^46,47^. Here we used a simple sampling strategy where the probabilities of codons at each position are based on the reference sequence. However, it is also possible to sample iteratively — by resampling after each accepted edit — or in parallel, by simultaneous infilling of multiple masked positions^32^. This sampling choice is especially relevant in light of our finding that model codon preferences depend on local and distal codon context. Thus, the practical utility of codon MLMs depends strongly on how easily their probability distributions are converted into sequence designs.

Our *in vitro* screen provides direct evidence that these design choices have functional consequences. The conservative editing strategies of EnCodon and CaLM — which modify fewer than 12% of codons — consistently outperformed the more aggressive recoding of CodonTransformer_AA (47% edited) and random synonymous substitution (65% edited). Both SEAP activity and RNA abundance declined with increasing codon Hamming distance from the reference (Spearman ρ = −0.73 and −0.55, respectively), and this relationship held uniformly across the gene rather than being driven by specific regions. These data suggest that preserving the native codon context, particularly at positions where the reference already uses well-adapted codons, is more beneficial than wholesale recoding — a principle that conservative MLM sampling naturally enforces.

Our interpretability analyses help to clarify how MLMs differ from simpler k-mer-based approaches. As expected, we found that MLM predictions are sensitive to multiple forms of sequence perturbation, indicating that these models integrate information across a broad sequence context in a way that frequency-based methods cannot. Interestingly, we observed a difference in the sensitivity of model predictions to local and distal perturbations. Specifically, perturbing a relatively small local window around a masked position caused substantial changes in model outputs, demonstrating that local context is necessary for predictions and is not redundant with distal context. However, restoring this local context within an otherwise perturbed sequence was not sufficient to recover the original predictions. This suggests that the models learn complex interactions between local and global sequence context, such that distal context modulates the output but cannot by itself restore it — a finding with direct implications for sequence design, as changing codons in one part of the sequence has complex effects on model outputs for another part. Altogether, our interpretability analyses reveal variable model sensitivity to synonymous, nonsynonymous, and masking perturbations at both local and distal scales, demonstrating that these MLMs incorporate broad sequence context when making predictions, differentiating themselves from frequency-based approaches for codon optimization.

Our study has several limitations. First, the benchmarking effort is not conclusive, in part due to the general lack of large-scale labelled datasets of synonymous coding sequences. Although some efforts in this direction exist^48^, the data are scarce and derived predominantly from bacteria rather than more therapeutically relevant systems such as human cell lines. The evaluation, training, and fine-tuning of codon MLMs would greatly benefit from expanded datasets of synonymous variants, both in terms of the number and diversity of proteins and the number of synonymous variants per protein. Second, we restricted our analysis to three prominent codon MLMs and we did not include supervised or autoregressive codon or nucleotide models^24,49,50^ in our benchmarks. This is due to the technical challenges of head-to-head comparisons across model classes, particularly when some may require fine-tuning on coding sequences. Future iterations should aim to incorporate these models. Third, our *in vitro* screen was restricted to a single protein (SEAP), and the generalizability of model rankings to other proteins, host organisms, or expression systems remains to be established. The screen also did not assess protein quality metrics such as glycosylation, folding, or secretion efficiency, which may be influenced by codon usage through effects on co-translational folding kinetics. Notably, the strong performance of conservative recodings in our screen may partly reflect the fact that the SEAP reference sequence is already well-adapted for human expression; for poorly expressed or non-human reference sequences, more aggressive recoding could prove beneficial.^35,51^

Beyond the limitations of our study, current models are also limited in their utility for designing DNA and RNA sequences for nucleic acid-based medicines in two important ways. First, with few exceptions^52,53^, models are specialized on individual genetic elements such as enhancers, UTRs, or coding sequences. However, these elements are known to interact, often nonlinearly, in their determination of molecular phenotype. For example, the RNA secondary structure surrounding the start codon is jointly determined by the 5′ UTR and the beginning of the coding sequence and can influence translational initiation — a clear case where modelling coding sequences in isolation may miss functionally relevant interactions. Moving beyond isolated coding sequence models to frameworks that capture interactions with regulatory elements could lead to more effective optimization strategies^52–55^.

Second, it may be important to model splicing jointly with coding sequences, as these processes interact in at least two ways. Depending on whether the construct contains an intronless or intron-containing transgene, the influence of GC content on expression may differ^30,56^. Because codon MLMs are trained on exonic sequences from diverse genes, including many intron-containing genes, the synonymous substitutions they propose may not be the same ones that maximize expression in intronless constructs. Additionally, MLM-suggested edits may inadvertently introduce cryptic splice sites, thereby reducing full-length protein expression. While our OpenSpliceAI analysis supports this concern in conventional methods – detecting predicted cryptic splice sites in one random variant, one commercial vendor variant, and the max. CAI construct – all MLM variants were free of such sites. This finding suggests that context-aware MLMs may implicitly learn to avoid splicing competent motifs. However, whether this effect is an inherent feature of their architecture or a byproduct of training data constraints requires further systematic validation.

Future work should aim to integrate these intertwined processes within a unified modelling framework. An ideal next-generation model would combine backtranslation-compatible generation, explicit conditioning on host species, control over unwanted motifs such as CpG dinucleotides, and conditional generation for target properties such as expression level. In practice, MLM-guided editing can be coupled with supervised predictors or explicit constraints to filter candidate sequences for specific therapeutic objectives^57,58^. Codon MLMs may therefore be most useful not as standalone optimizers but as generators of diverse, natural-like candidates for downstream filtering and selection - an approach validated by our screen, where sampling multiple variants from each model and selecting the best performers proved more effective than relying on a single design. Developing such integrated approaches will be an important step toward making sequence language models directly actionable for the rational design of nucleic acid-based medicines.

## Methods

### Selection of diverse proteins

To place our characterization of model-generated coding sequences and our interpretability analyses in a therapeutically relevant setting, we curated a dataset of human proteins matched to their coding DNA sequences and protein family annotations. We began with the CaLM training set, which provides a large collection of curated coding sequences indexed by ENA identifiers, but does not include species annotations or protein family information. To restrict this resource to human sequences, we queried NCBI for each entry and retained only records annotated as *Homo sapiens*.

We then obtained protein family annotations from UniProt by downloading all reviewed human Swiss-Prot entries, which include UniProt protein names (for example, 1433B_HUMAN), amino acid sequences, and curated family annotations. Using the UniProt ID mapping file, we mapped the human sequences from the CaLM-derived set to UniProt entries. To ensure that these mappings were correct, we translated each coding sequence and aligned the resulting amino acid sequence to the corresponding UniProt protein sequence, retaining only matches with an alignment score greater than 0.99. This procedure yielded 6,184 entries with verified coding sequences and protein family annotations.

To preserve diversity across protein families while limiting overrepresentation of large families, we randomly subsampled families containing more than 50 entries to 50 sequences and retained all families represented by more than one sequence otherwise. Finally, to simplify downstream analyses and ensure compatibility with model input constraints, we restricted the dataset to proteins between 300 and 1,023 amino acids in length, resulting in a final set of 2,012 sequences.

To compare in silico samples across a set of representative and diverse proteins, we selected five proteins by choosing one random member from each of the five largest protein families represented in the set of 2,012 proteins. The selected proteins were SLC25A3 (Q00325), CYP24A1 (Q07973), TRIM9 (Q9C026), TSP50 (Q9UI38), and ADMR (O15218).

### Benchmarking datasets

We used the following datasets in our benchmark, filtering out any sequences that were non-coding:

- Mean Ribosomal Load - curated by Fradkin *et al*.^59^, originally from Sugimoto and Ratcliffe^60^. After filtering for sequences with known CDS coordinates, the dataset contained 9,933 sequences. Note that this dataset is derived from a single cell type - kidney cancer cells - potentially leading to more noise as compared to other datasets in the benchmarking panel.
- Protein and transcript abundance - curated by Outeiral & Deane^20^, originally collected from PAXdb^61^ repository and Human Protein Atlas^62^ with 11,790 and 5,485 sequences, respectively. The data represent averaged values across different human tissues.
- mRFP abundance - curated by Li et al.^27^ and originally derived from Nieuwkoop et al.^4^ for mRFP expression in *E.coli* from 1,455 unique synonymous sequence variants.
- MLOS flu vaccines - generated by Li et al.^27^ measuring hemagglutinin antigen expression relevant for flu vaccines and comprising 167 sequences.^43^
- mRNA half-life: curated by Fradkin et al.^59^ and originally assembled by Agarwal and Kelley^43^ with each data point being derived from PCA (first principal component) of a range of different mRNA measurements for the same gene. Overall, there were 12,969 sequences in this dataset.
- Nanoluc abundance: generated by Leppek et al., 2022 the dataset is composed of 24 synonymous Nanoluc luciferase sequences and associated abundance readouts from transfected HEK293T cells after 24 hours.
- GFP fluorescence: generated by Mordstein et al.^30^ the dataset is composed of GFP fluorescence FACS sorting readouts under three conditions - (i) HeLA with introns (182 sequences) (ii) HeLA without introns (184 sequences) (iii) HEK293 without introns (185 sequences).

### Linear probing

To evaluate model performance, we froze the weights of the MLMs and linearly regressed the molecular phenotypes of our benchmark on the sequence embeddings. To do so, we performed a forward pass on each sequence and extracted its embedding from the final layer of each model. CodonTransormer requires a species tag as input, which we set to ‘Homo sapiens’ for all tasks in our benchmark except mRFP abundance, for which we set the tag to ‘Escherichia coli general’. For each fold in our 5-fold cross-validation, we used the training set to normalize the embeddings with StandardScaler from scikit learn and fit these with PCA, with number of features set to 320 (or no reduction if the number of features is lower than that) and transformed both the training and test sets. We then trained an elastic net with a maximum of 5,000 iterations to predict the labels from the transformed embeddings of the training set, and used the resulting model to predict the labels from the transformed embeddings of the test set, recording the Pearson correlation coefficient. For our baselines, we performed the procedure above using k-mer frequencies instead of model embeddings. Additionally, we defined the concatenation of all MLM embeddings as the combined MLM features and the concatenation of k-mers and MLM embeddings as the k-mers + MLM. Note that the number of features is still capped at 320 despite the larger number of initial set after concatenation.

### Codon preference

Codon preference was calculated by normalizing synonymous codon logits and computing the entropy of this distribution using the number of synonyms as the base of the logarithm, which facilitates comparison across amino acids with different numbers of codons. Entropy was then subtracted from 1 to make the values more intuitive, such that low and high values correspond to low and high biases in codon preference, respectively.

### Choosing codons to mask

For CaLM and EnCodon,we selected codon positions to mask based on amino acid fidelity. Specifically, we chose codon positions where amino acid fidelity exceeded an arbitrary threshold of 0.8, and that were at least 100 codons away from the beginning and end of the sequence. Additionally, we subsampled over-represented amino acids such that none appears more than 700 times in the final set. For CodonTransformer, such selection is not necessary, because the model’s tokenization scheme yields non-zero output probabilities only at the input amino acids’s synonymous codons. To provide a fair comparison of the three models, we used the masked positions from CaLM and EnCodon for CodonTransformer.

### Local and distal context perturbation

To perturb local sequence context, we corrupted the flanking sequence on both sides of each masked position using window sizes ranging from 1 to 100. The smallest window size of 1 means that only the codons immediately flanking the masked position are perturbed, whereas the largest window size of 100 means that 100 codons to either side of the masked position are perturbed. In contrast, to perturb distal sequence context, we corrupted the entire sequence, and incrementally restored the sequence around the masked position using window sizes ranging from 0 to 500, in increments of 5. The smallest window size of 0 means that all codons on either side of the masked position are perturbed, whereas the largest window size of 500 means that all codons beyond the window are perturbed.

For each masked position at each window size, we ran a forward pass on the MLM using the perturbed sequence to obtain codon probabilities at the masked position, which we used to compute amino acid fidelity and codon preference. For synonymous and nonsynonymous perturbations, we repeated this process 10 times, perturbing the sequence using randomly chosen synonymous and nonsynonymous codons, respectively. In our analysis of asymmetric perturbations, we followed the same procedure, but only perturbed the sequence on one side of the masked position.

### Vector Design

Codon-optimized SEAP variants were synthesized as gene fragments (Twist Bioscience) and directionally assembled into the donor vector via Golden Gate assembly. The reporter donor vector utilizes a dual-promoter configuration driving a selection marker and the SEAP reporter cassette. Assemblies were performed according to the NEBridge protocol (New England Biolabs). Sequence validation was performed by nanopore sequencing (Microsynth).

### Cell Culture

Engineered HEK293A cells were maintained as adherent monolayers in standard high-glucose DMEM supplemented with 10% FBS, L-glutamine, penicillin-streptomycin, and selection antibiotics at 37 °C in 5% CO₂. Cells were seeded at 2.5 × 10⁶ cells per T75 flask in 12 mL medium and maintained below 80% confluency.

### Transfections and Stable Cell Line Generation

Cells from a homogeneous batch were seeded at 11,000 cells per well in a 96-well plate 16 h prior to transfection. Site-specific genomic integration was performed using a recombinase-mediated landing pad system. Cells were co-transfected with a recombinase expression plasmid and donor plasmid using commercial transfection reagents according to standard protocol. After 48 h, 100 µl conditioned supernatant was collected for transient SEAP measurements. Stable polyclonal populations were selected with 0.2 mg/ml zeocin (Gibco).

### SEAP Reporter Assay

SEAP activity in cell culture supernatants was quantified by kinetic colorimetric assay adapted from Schlatter *et al*.^63^. Supernatant (100–200 µL) was heat-inactivated at 65 °C for 30 min. Twenty microlitres of heat-inactivated supernatant was combined with 180 µL assay reagent. Hydrolysis of pNPP to 4-nitrophenol was monitored at 405 nm at 2-min intervals over a 16-min linear window at 37 °C using a commercial plate reader.

### DNA/RNA Extraction and High-Precision Transcript Quantification via dPCR

Immediately after supernatant collection for stably integrated SEAP measurements, DNA and RNA were co-extracted using the AllPrep 96 DNA/RNA Kit (QIAGEN) according to the manufacturer’s protocol. Nucleic acid concentrations were determined using a Lunatic spectrophotometer (Unchained Labs). cDNA was synthesized using the Transcriptor First Strand cDNA Synthesis Kit (Roche) with anchored oligo(dT) primers. RNA was quantified by digital PCR (QIAcuity Eight, QIAGEN) using human *HPRT*1 as a reference gene (4333768, Thermo Fisher Scientific) and HEX-labeled SEAP-specific primers. Genomic DNA copy number was determined using the TaqMan Copy Number Reference Assay for human RNase P (4403328, Thermo Fisher Scientific).

### Data Normalization and Statistics

Transient SEAP activity was normalized to the reference variant and expressed as Z-scores across biological replicates. Stably integrated SEAP activity was normalized to genomic DNA copy number (diploid equivalents) to account for variation in integration efficiency across wells. RNA levels were normalized to *HPRT*1 expression. Specific productivity was calculated as the ratio of relative stably integrated SEAP activity to relative RNA abundance.

All statistical analyses were performed in Python 3.12.3 using SciPy (v1.17.1), statsmodels (v0.14.6), NumPy (v2.4.6), and pandas (v3.0.3). Group comparisons (MLMs, Random, Reference) were performed using Tukey’s HSD post hoc test for groups with n ≥ 2. For comparisons involving the Reference (n = 1), one-sample t-tests with Bonferroni correction for the number of single-group comparisons were used. Correlations were assessed using Spearman’s rank correlation coefficient (ρ) and Pearson’s R². Linear regression was performed using ordinary least squares, with 95% confidence intervals computed from the standard error of the fit. Pairwise sequence distances were calculated as codon Hamming distance (fraction of differing codons out of 507). All tests were two-sided. Data are presented as mean ± s.d. from n = 3 biological replicates unless otherwise stated. Figures were generated using Matplotlib (v3.10.9).

### Splicing predictions

Cryptic splice sites in the SEAP variants were predicted using OpenSpliceAI (v0.0.4)^44,45^. Predictions were made using the human OSAI-MANE models trained on GRCh38/MANE transcripts with a 10,000 nt context window. We reproduced the standard workflow by using the ensemble of the five independently trained models provided by the authors. Each full-length transcript, including its 5’ UTR, was one-hot-encoded and split into overlapping windows. The per-position acceptor and donor probabilities from the five models in the ensemble were averaged to yield a single ensembled prediction per nucleotide. For each variant, a cryptic in-CDS acceptor score was defined as the maximum predicted acceptor probability at any position within the coding sequence.

## Acknowledgments

We thank Dr. Grzegorz Kudla for generously sharing a processed version of the dataset described in Mordenstein, *et al*.^30^. The authors are grateful to Markus Grosch for sharing the engineered HEK293A cell line with the incorporated landing pad site.

## Contributions

S.T. performed all machine learning analyses. K.S. and F.N. designed the *in vitro* screen. K.S. performed, analyzed and visualized the *in vitro* experiments. T.C., J.L.P., S.A., J.S., S.T. and C.N. interpreted the results and helped improve the manuscript. S.T. and K.S. wrote the manuscript with input from all authors. C.D.D. performed the splicing analysis. T.C. and J.L.P. acquired funding. S.A., T.C. and J.L.P. supervised the project.

## Supplementary Figures

**Figure S1:**
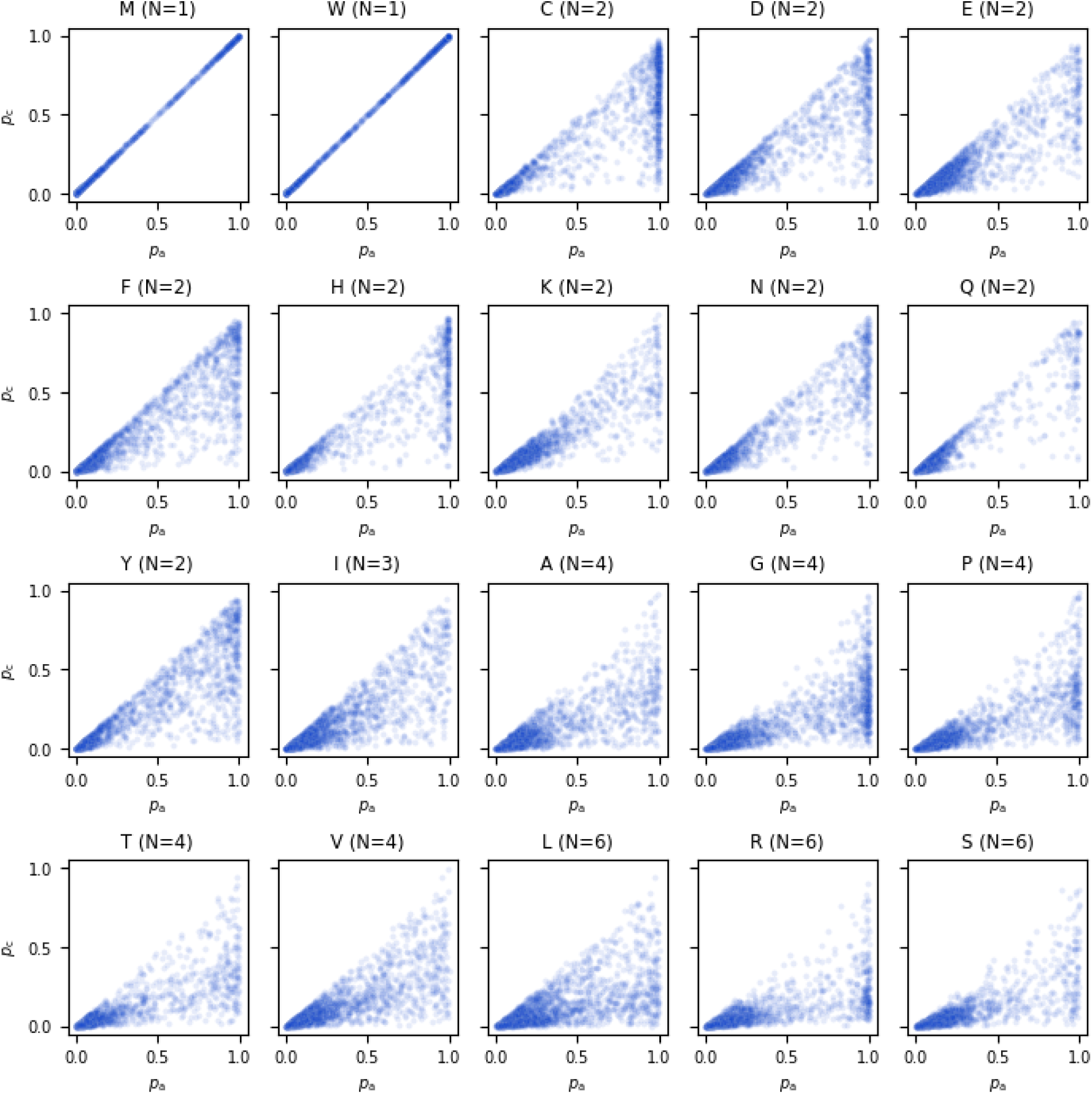
The relationship between reference codon probability and amino acid fidelity varies across amino acids. Scatter plot of *p*_c_ vs *p*_a_ per masked codon in each of 2,012 proteins, stratified by amino acid. The title of each subplot indicates the amino acid and its number of codons. Data pertain to model predictions from CaLM.

**Figure S2:**
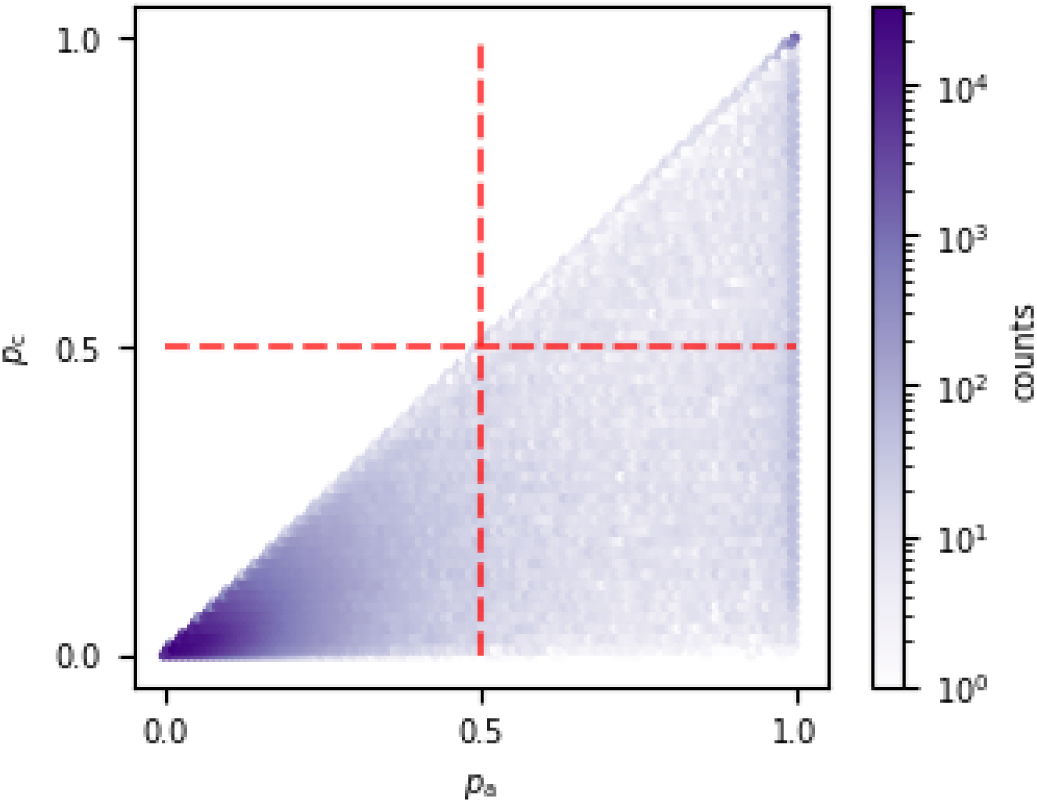
Density plot of *p*_c_ vs *p*_a_ per codon in each of 2,012 proteins using EnCodon (n=1,087,855 masked codons).

**Figure S3:**
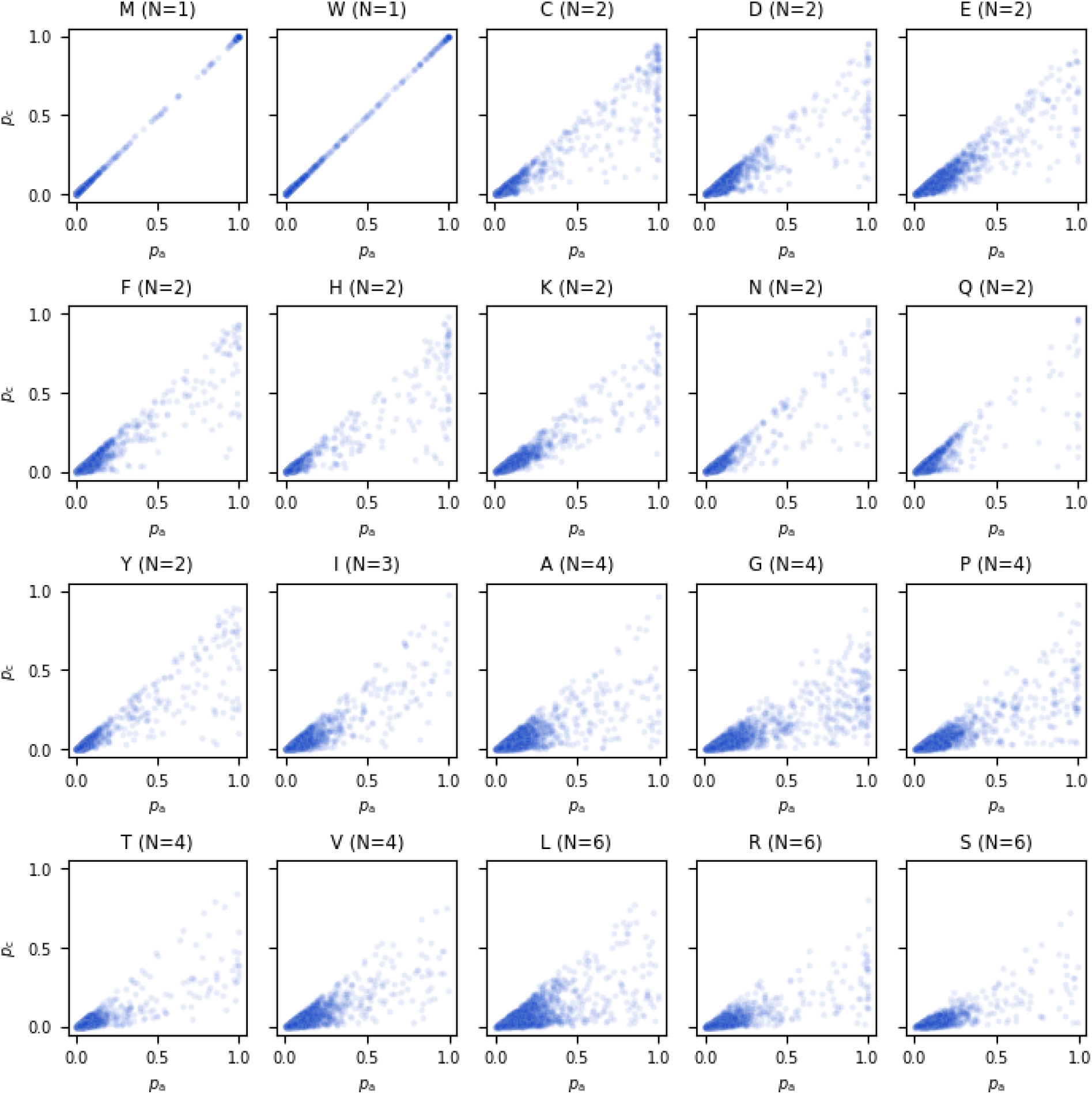
The relationship between reference codon probability and amino acid fidelity varies across amino acids. Scatter plot of p_c_ vs p_a_ per masked codon in each of 2,012 proteins, stratified by amino acid. The title of each subplot indicates the amino acid and its number of codons. Data pertain to model predictions from EnCodon.

**Figure S4:**
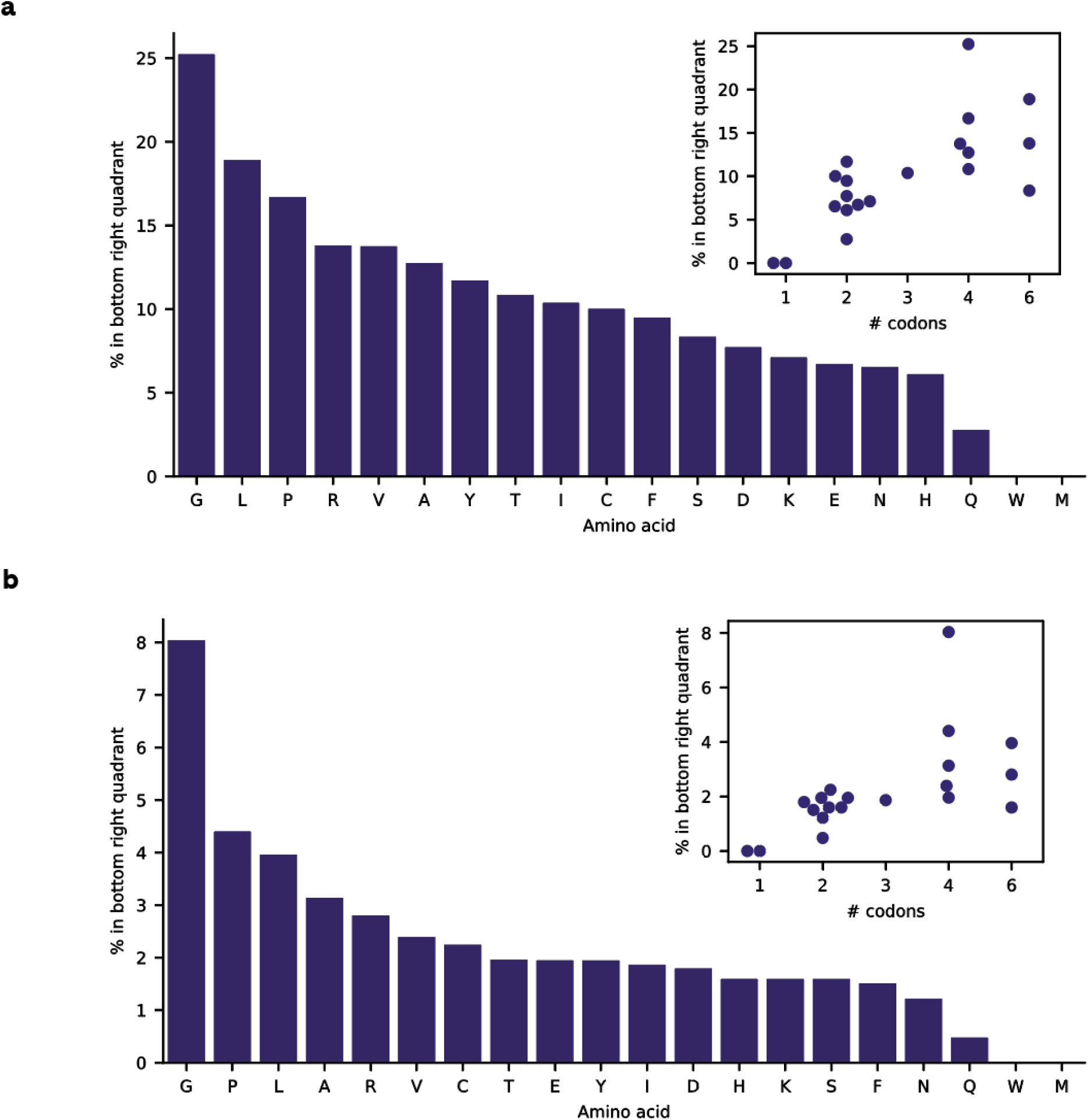
The percentage of positions amenable to synonymous recoding varies across amino acids. Bar plots of the number of masked positions that fall in the bottom right quadrant of Fig. 1b, stratified by amino acid, for (a) CaLM and (b) EnCodon. The insets show the number of masked positions in relation to the number of codons per amino acid.

**Figure S5:**
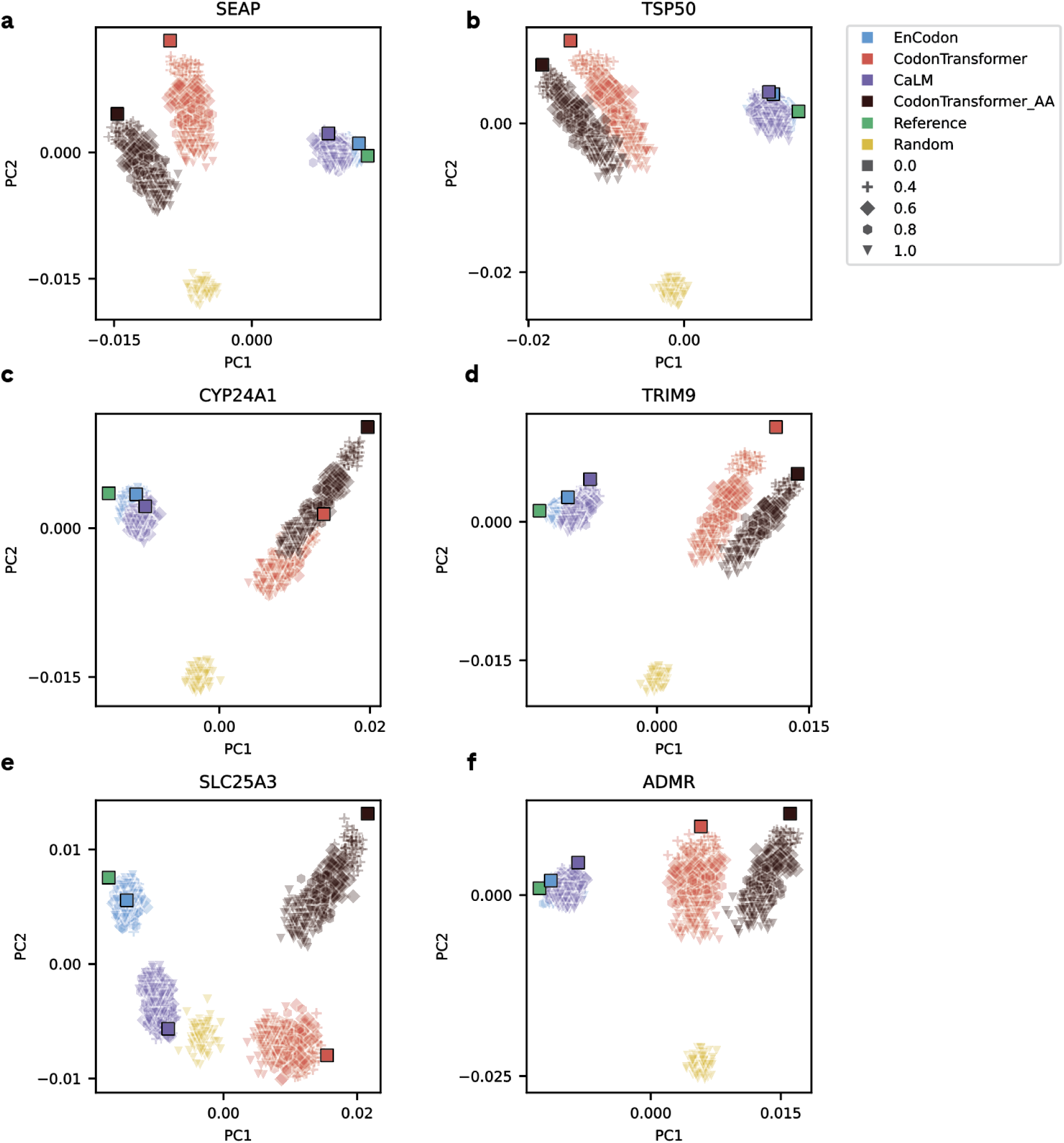
Sampled synonymous sequence variants differ in their statistical signatures, as evidenced by their separation in principal component space. Each panel pertains to one of six human proteins: (a) SEAP, (b) TSP50, (c) CYP24A1, (d) TRIM9, (e) SLC25A3, (f) ADMR. Symbol colors correspond to models and shapes to temperatures (legend). Darker symbols indicate sequences derived deterministically, by sampling the highest-probability codon at each masked position.

**Figure S6.**
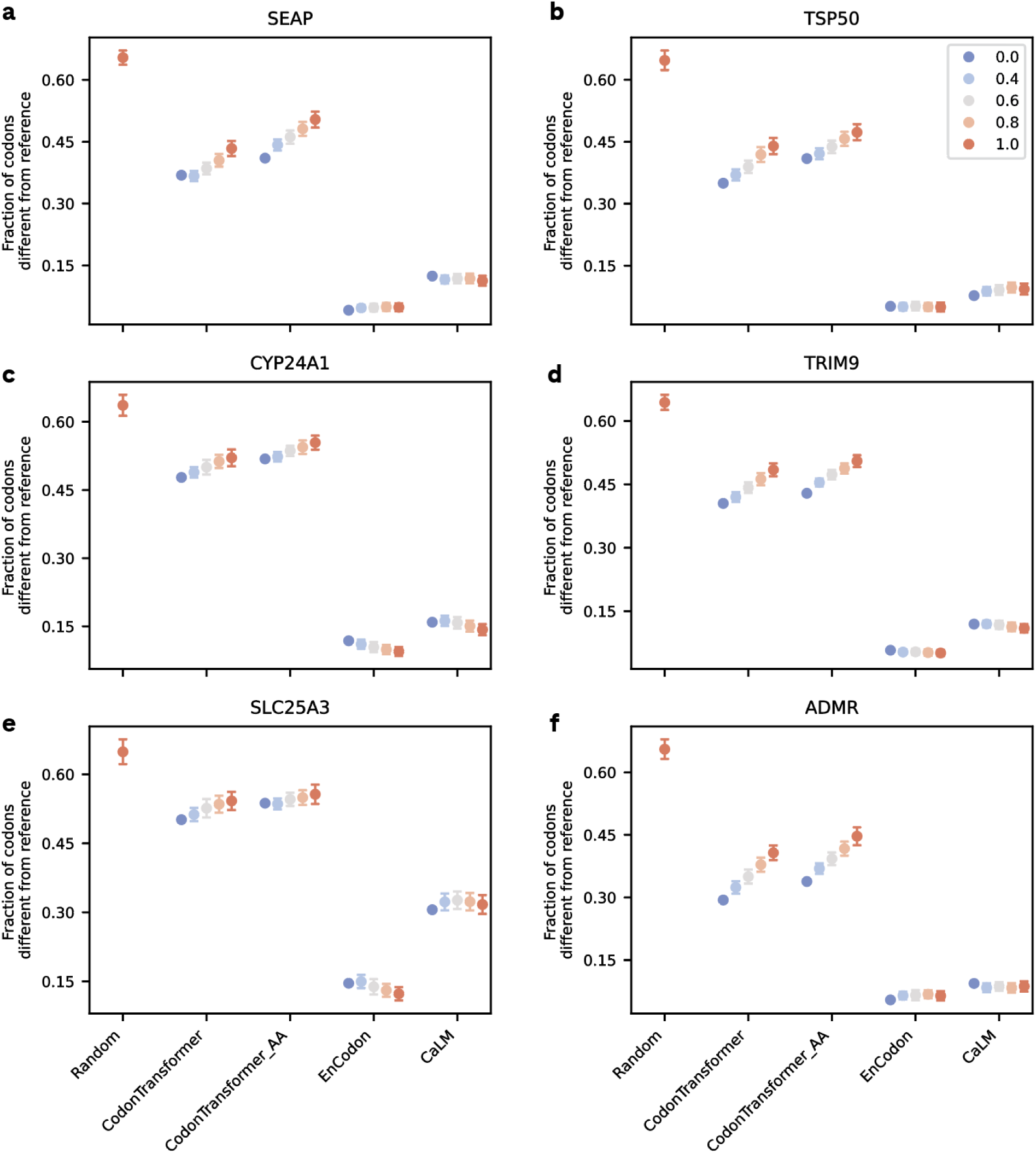
Sampled synonymous variants differ in their edit distance (in codons) from the reference sequence across models and temperatures (legend). Each panel pertains to one of six human proteins: (a) SEAP, (b) TSP50, (c) CYP24A1, (d) TRIM9, (e) SLC25A3, (f) ADMR. Symbols represent the mean and error bars the standard deviation across sampled variants. Symbol colors correspond to temperature (legend).

**Figure S7.**
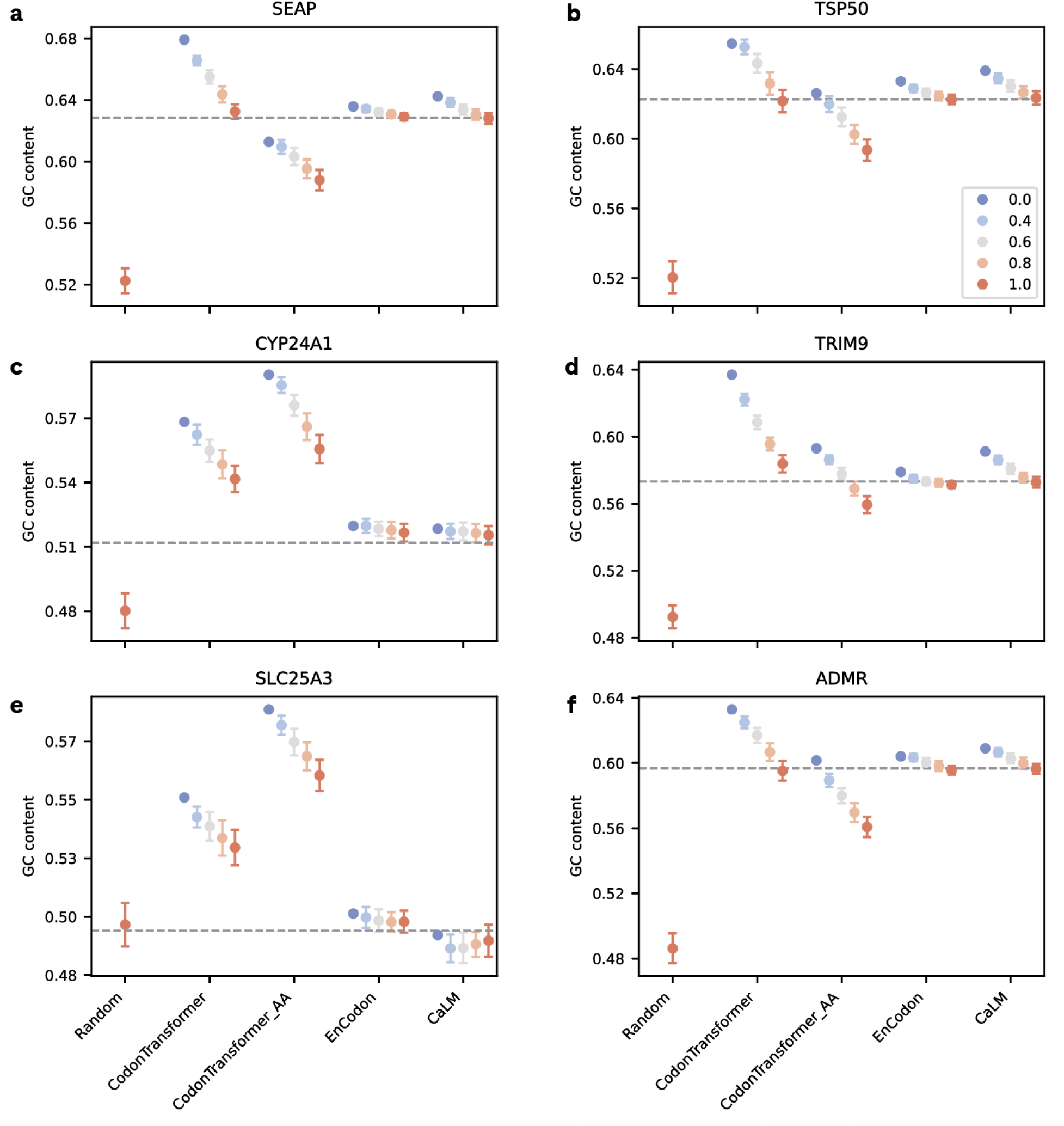
Sampled synonymous sequence variants differ in their GC content across models and temperatures (legend). Each panel pertains to one of six human proteins: (a) SEAP, (b) TSP50, (c) CYP24A1, (d) TRIM9, (e) SLC25A3, (f) ADMR. Symbols represent the mean and error bars the standard deviation across sampled variants. Symbol colors correspond to temperature (legend). The dashed horizontal line indicates the GC content of the reference sequence.

**Figure S8.**
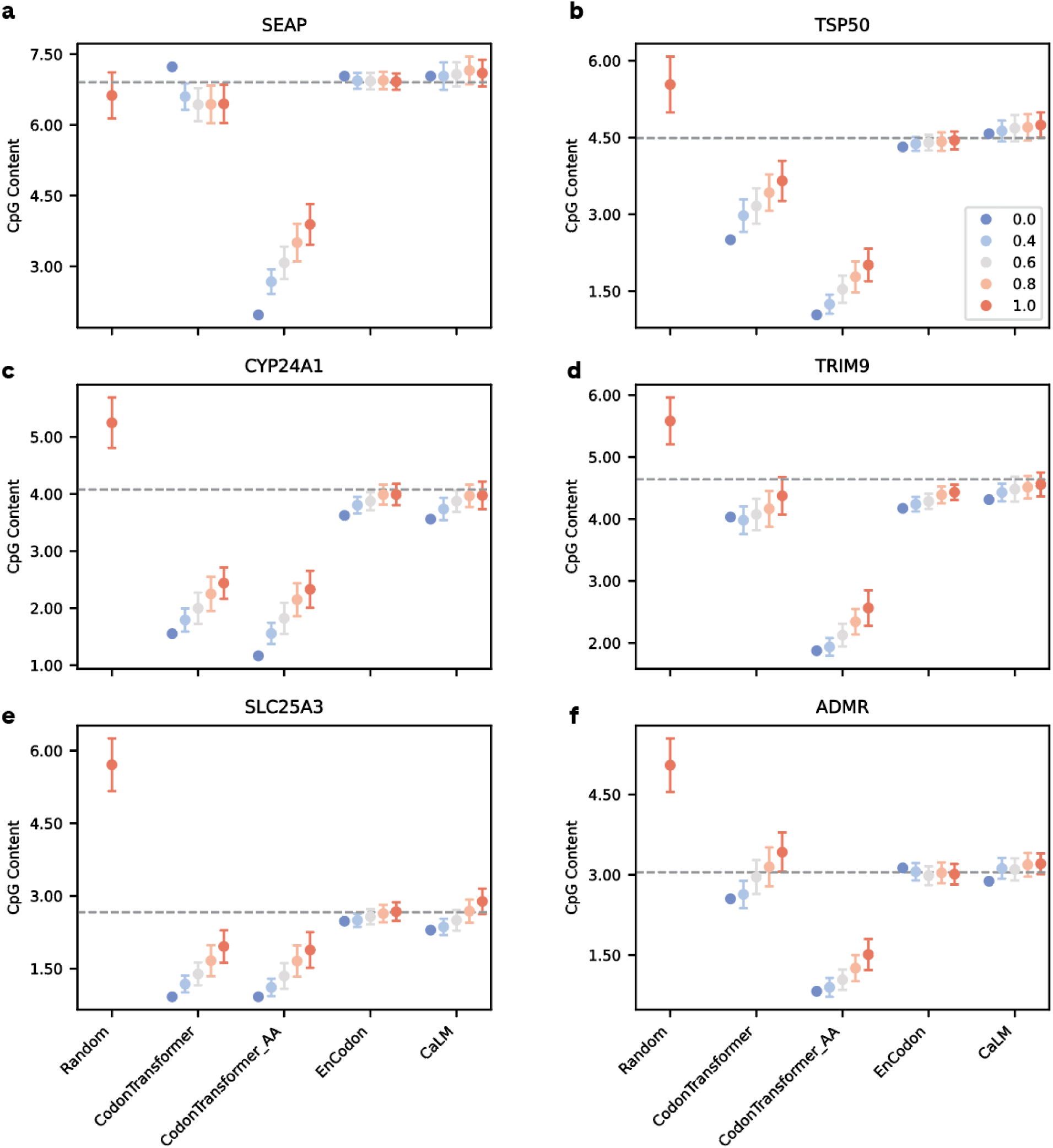
Sampled synonymous sequence variants differ in their CpG dinucleotide content, shown as a percentage of sequence length, across models and temperatures (legend). Each panel pertains to one of six human proteins: (a) SEAP, (b) TSP50, (c) CYP24A1, (d) TRIM9, (e) SLC25A3, (f) ADMR. Symbols represent the mean and error bars the standard deviation across sampled variants. Symbol colors correspond to temperature (legend). The dashed horizontal line indicates the CpG content of the reference sequence.

**Figure S9.**
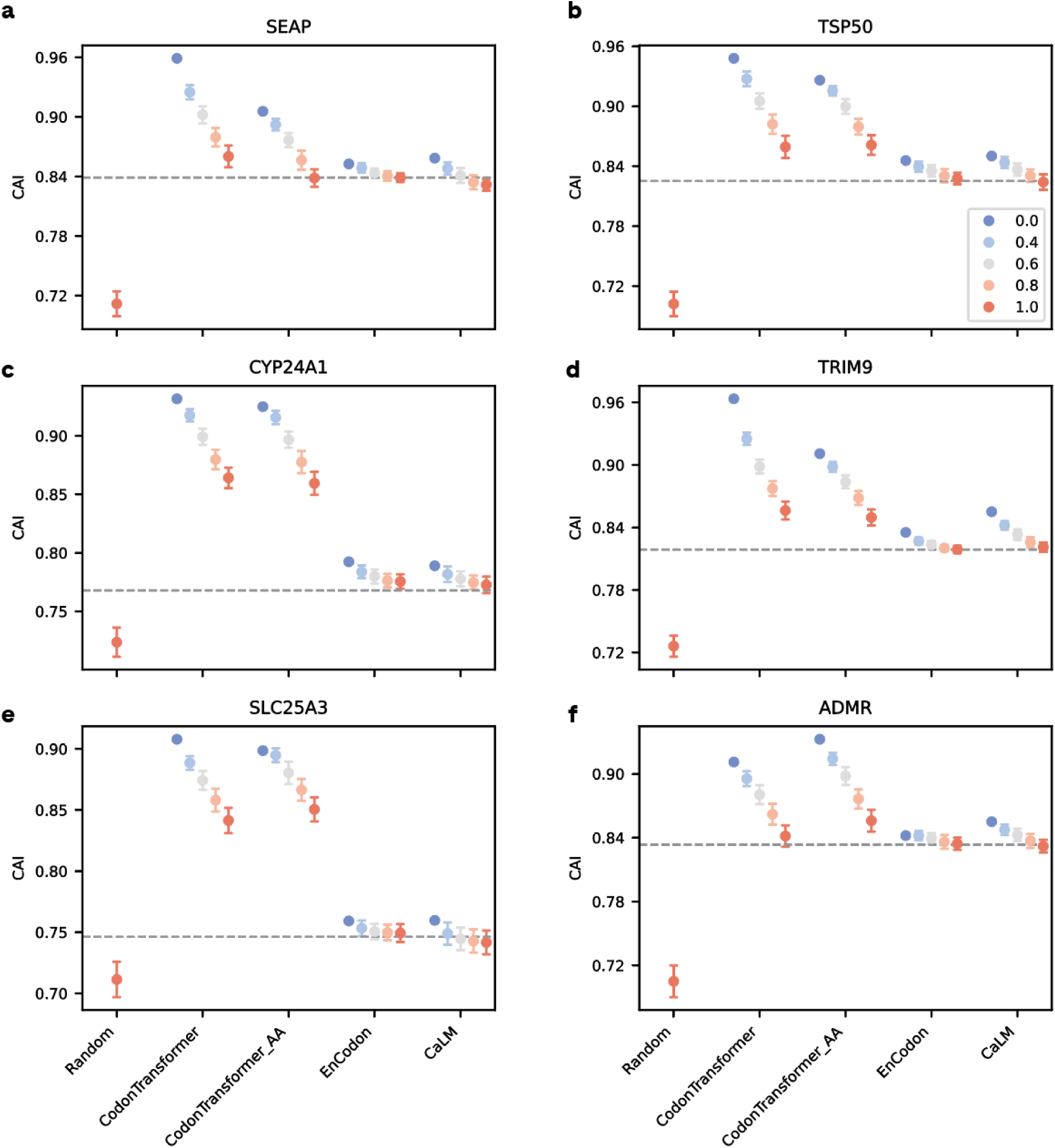
Sampled synonymous sequence variants differ in their CAI values across models and temperatures (legend). Each panel pertains to one of six human proteins: (a) SEAP, (b) TSP50, (c) CYP24A1, (d) TRIM9, (e) SLC25A3, (f) ADMR. Symbols represent the mean and error bars the standard deviation across sampled variants. Symbol colors correspond to temperature (legend). The dashed horizontal line indicates the CAI of the reference sequence.

**Figure S10:**
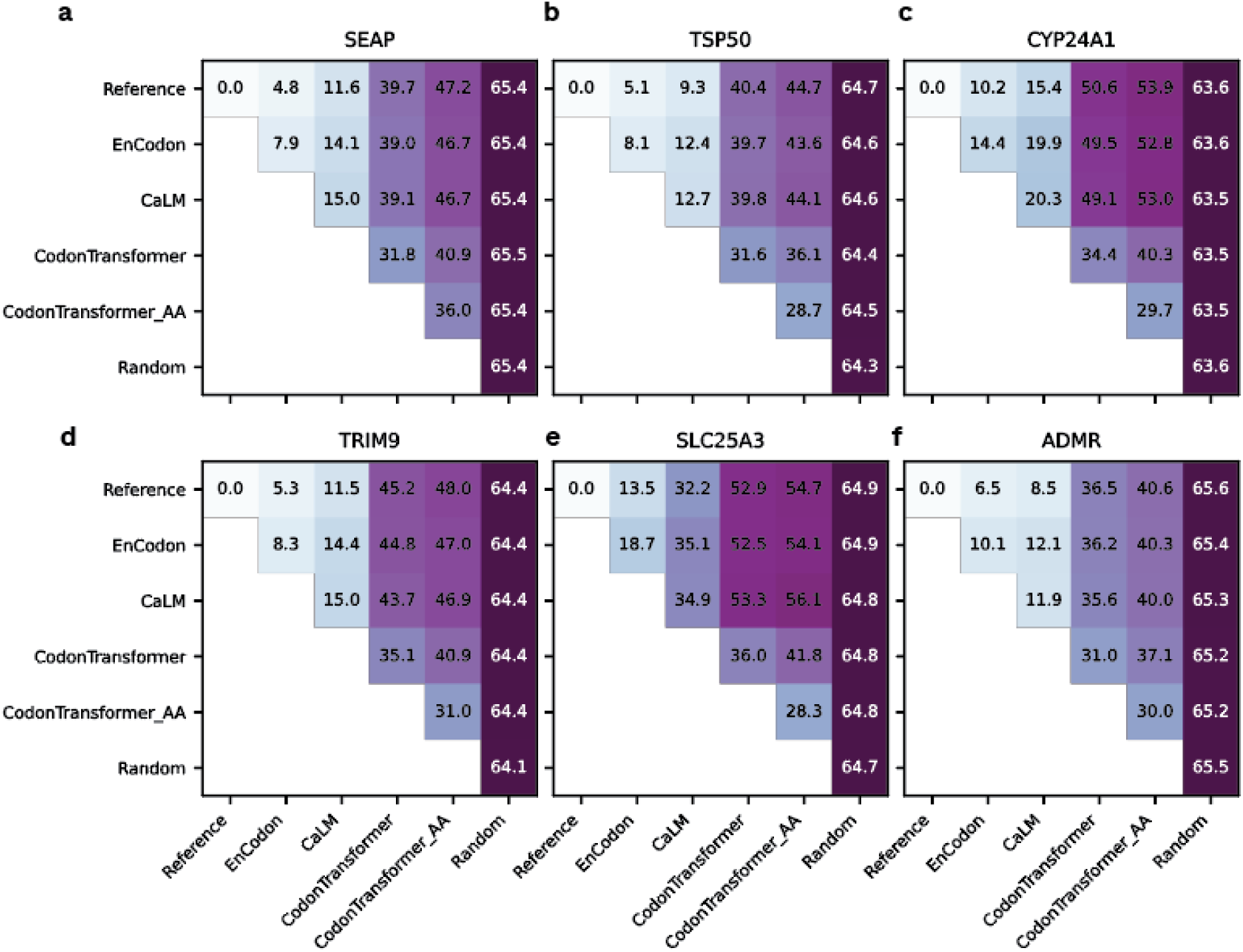
Sequence diversity within and between models. Each panel pertains to one of six human proteins: (a) SEAP, (b) TSP50, (c) CYP24A1, (d) TRIM9, (e) SLC25A3, (f) ADMR. Each matrix element shows the average pairwise difference, reported in percentage of codons, between synonymous sequence variants sampled from two models, which are indicated by the row and column names. The main diagonal shows within-group pairwise differences.

**Figure S11:**
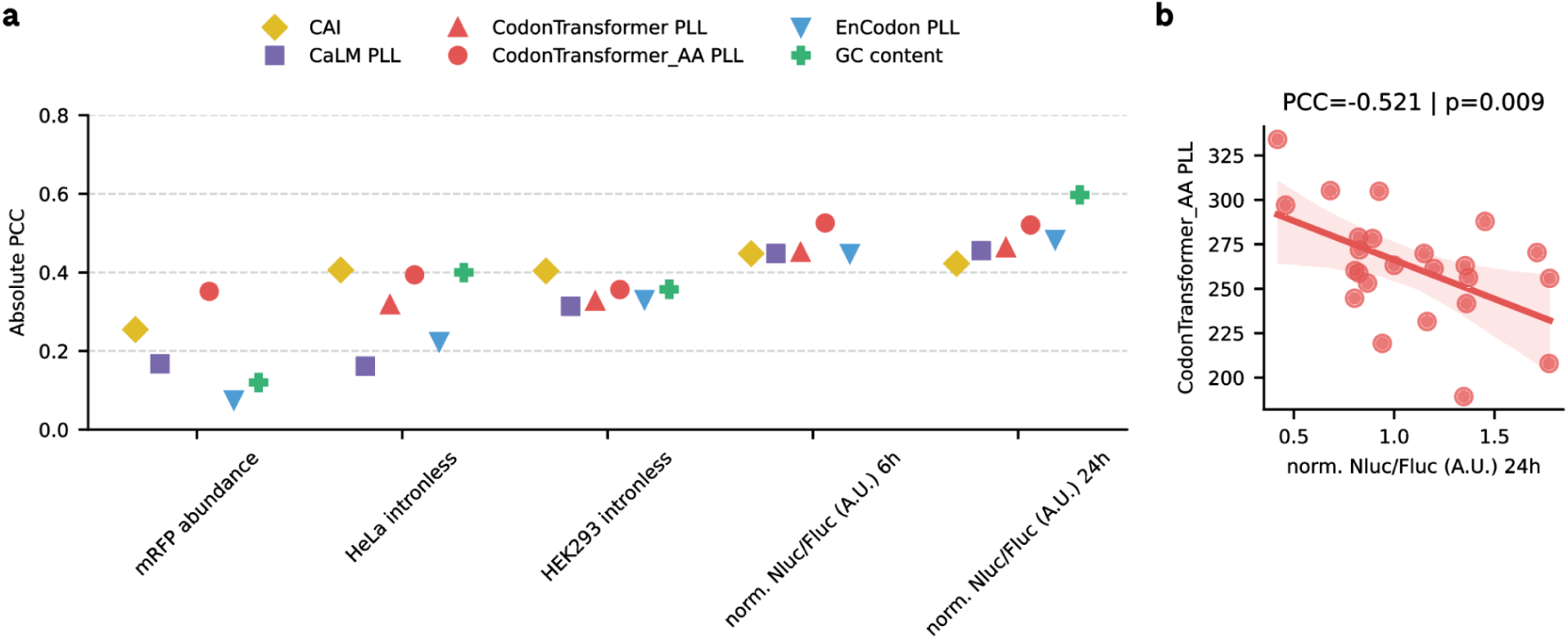
Pseudo log likelihoods (PLLs) as zero-shot predictors of molecular phenotype for datasets of synonymous sequence variants, which are typically too small for fine-tuning. a, The absolute Pearson correlation coefficient (PCC) across datasets and models (see legend). The absence of a symbol indicates statistical insignificance (*p* > 0.05) for that dataset. b, Scatter plot of the PLL from CodonTransformer_AA and a proxy of translational efficiency (Methods), shown as a representative example.

**Figure S12:**
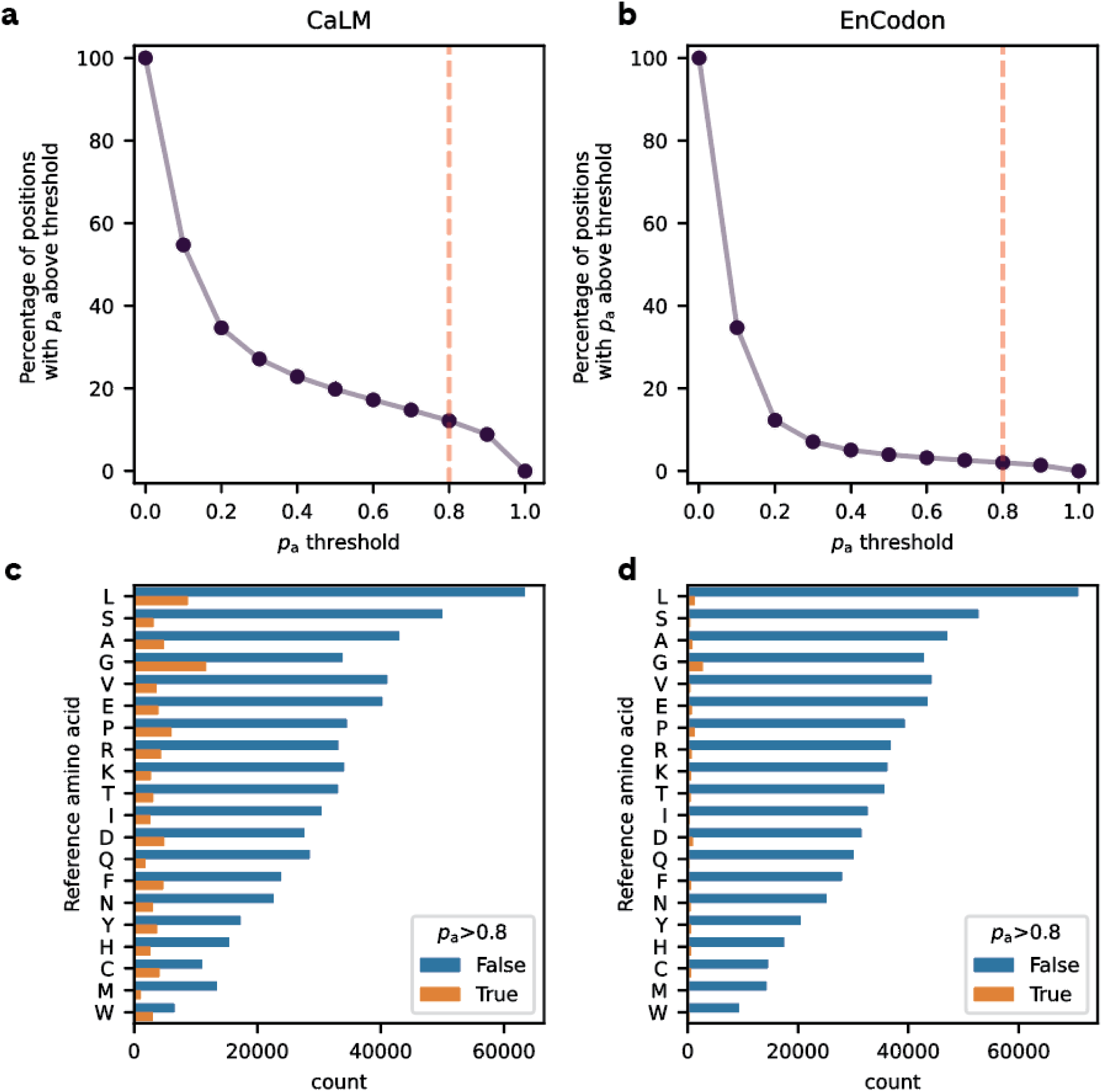
Choosing positions to mask. (a,b) Percentage of positions available for masking, shown in relation to the threshold on amino acid fidelity, *p*_a_, for CaLM and EnCodon (n=685,933). The dashed vertical line corresponds to the threshold used in our main text. (c, d) Counts of codon positions that do or do not exceed the threshold of *p*_a_ = 0.8, stratified by amino acid.

**Figure S13:**
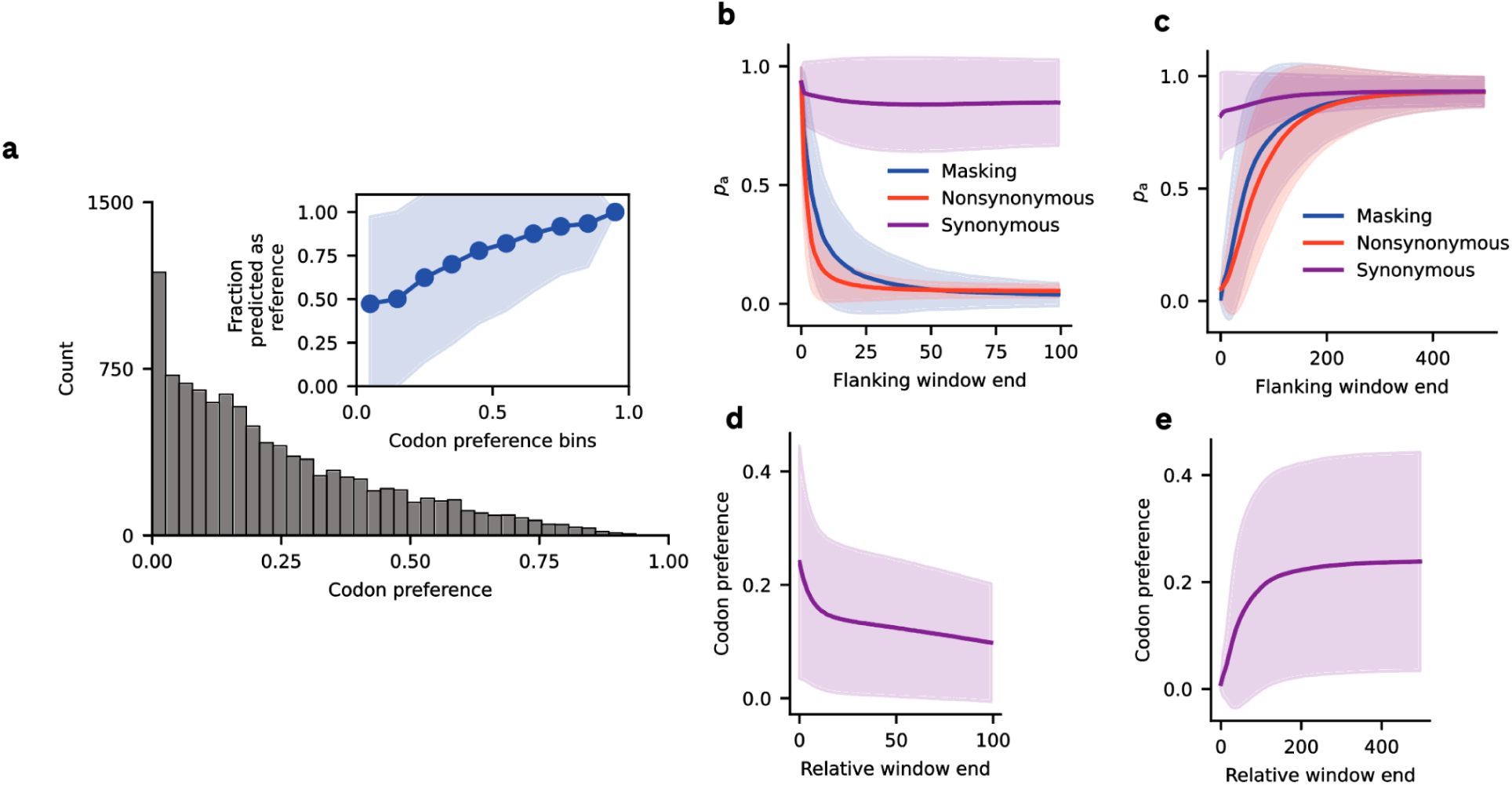
Local and distal context perturbations for EnCodon. a, Codon preference distribution in unperturbed sequences. The histogram shows the distribution of codon preference values for 10,132 codon positions where the input codon has a synonym. The inset shows the mean fraction of positions where the codon with the predicted highest probability corresponds to the input codon. b, Local context perturbations. The line plot shows amino acid fidelity, *p*_a_, in relation to the window size of the perturbation, stratified by perturbation type. c, Distal context perturbation. The line plot shows mean *p*_a_ across perturbation window sizes, stratified by perturbation type; d, The effect of local synonymous perturbation on codon preference. The line plot shows the change in codon preference with an increasing synonymous perturbation window. e, The effect of distal synonymous perturbation on codon preference. The line plot shows codon preference in relation to the window size of synonymous perturbation. In panels b-d, solid lines represent the mean, and shaded areas the standard deviation, across codon positions. Panels b and c pertain to data from all 10,577 sequence positions selected such that *p*_a_ is above 0.8 and d and e to 10,132 sequence positions. Note that for local perturbations (panels b, d), the window end refers to the perturbation area, whereas for distal perturbations (panels c, e), the window end refers to the unperturbed area.

**Figure S14:**
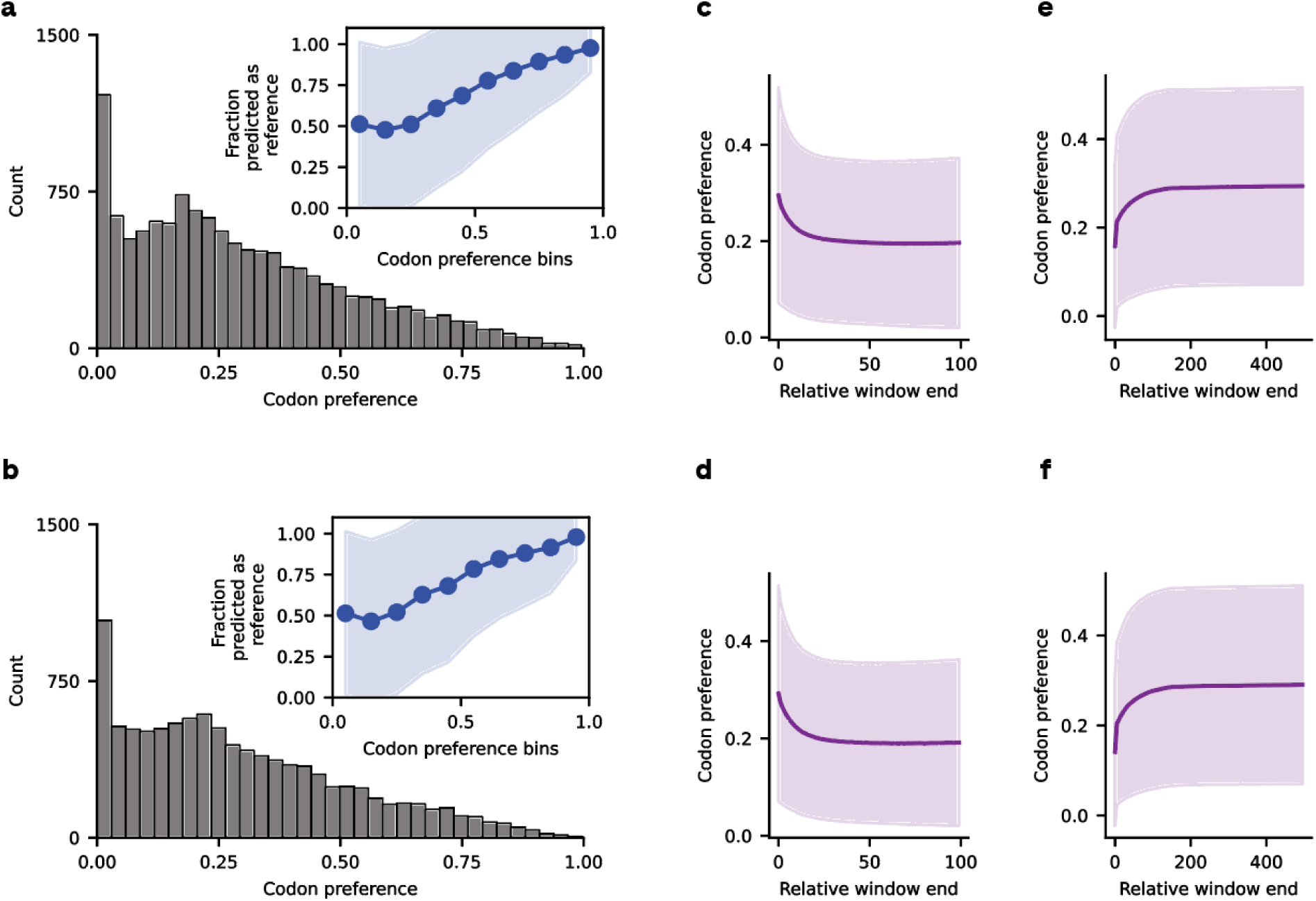
Local and distal context perturbations for CodonTransformer, using matched CaLM and EnCodon masked codons. a,b, Codon preference distributions for unperturbed sequences. Histograms show values for 12,600 and 10,132 masked codons corresponding to the positions analysed for CaLM and EnCodon, respectively. Insets show the mean fraction of positions at which the highest-probability codon matches the input codon; shading indicates the standard deviation. c, d, Local context perturbation using the CaLM and EnCodon masked codons, respectively. e, f Distal context perturbation using the CaLM and EnCodon masked codons, respectively. The line plots show the mean codon preference in relation to the window size of the perturbation.

**Figure S15:**
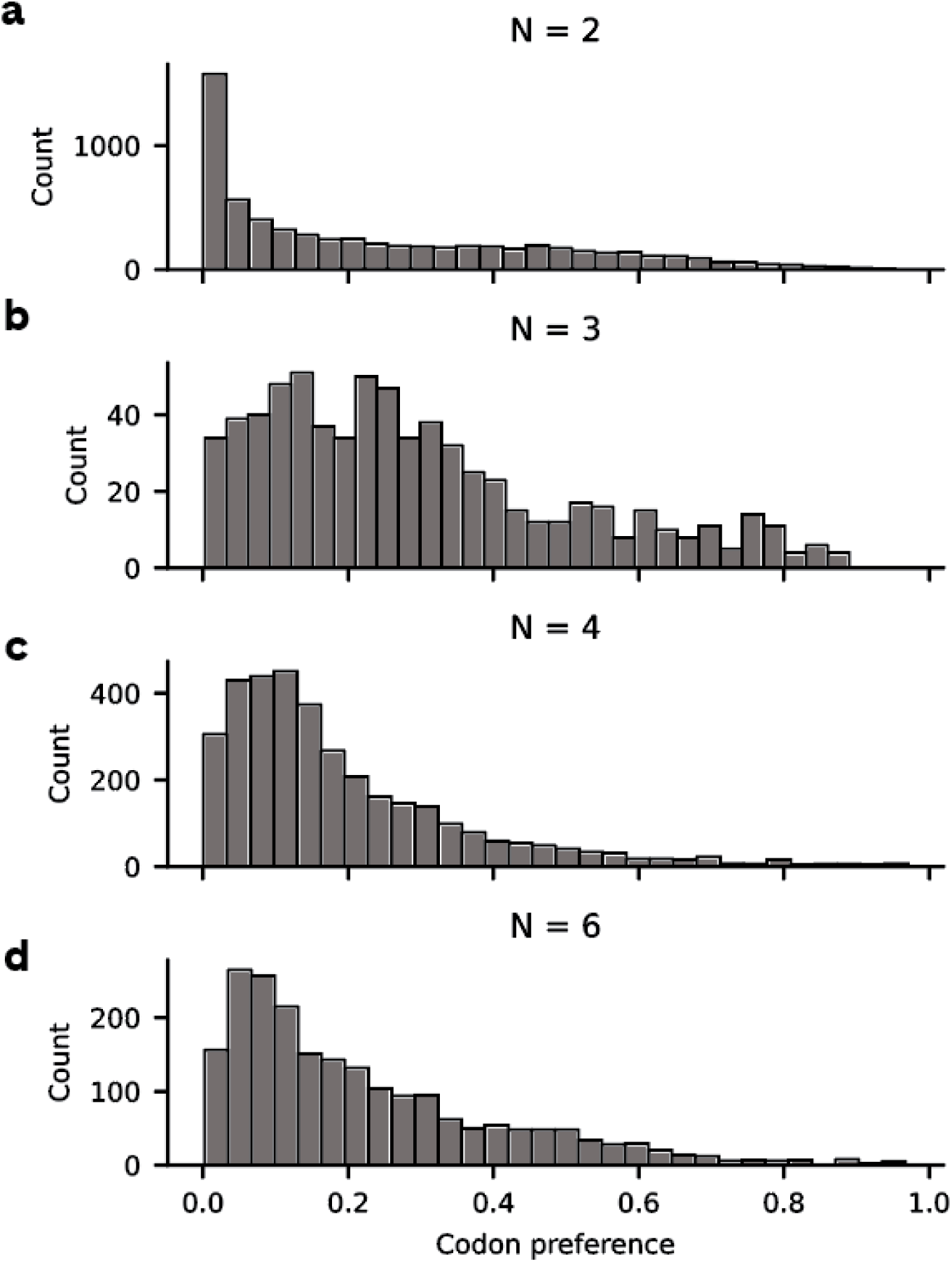
Codon preference distribution in unperturbed sequences, stratified by the number of codons per amino acid: (a) N = 2, (b) N = 3, (c) N = 4, and (d) N = 6. Data pertain to CaLM.

**Figure S16:**
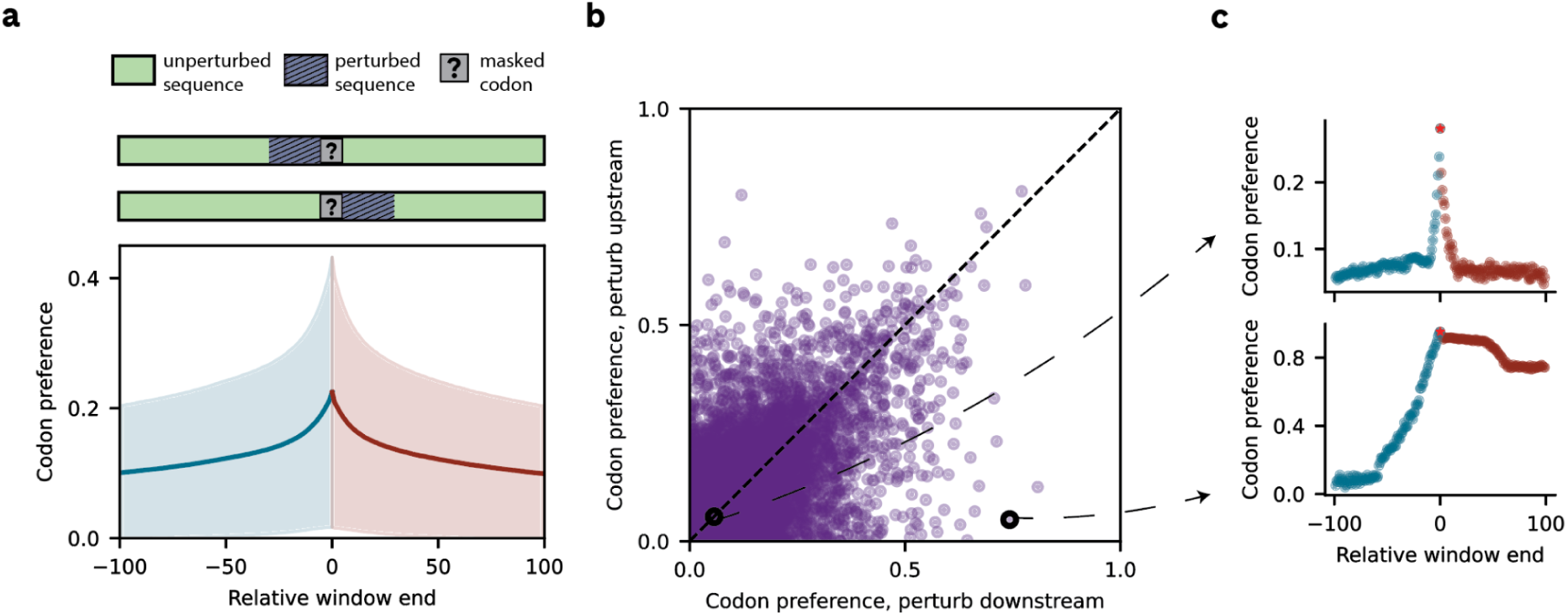
Model sensitivity to asymmetric, local context perturbation. a, The top diagram shows a schematic representation of the analysis. The bottom panel shows the average (solid line) and standard deviation (shaded area) of codon preference in relation to window size for asymmetric perturbation, where the sequence upstream (purple) or downstream (orange) of the masked position is perturbed. b, Comparison of codon preference when the sequence upstream and downstream of a masked position is perturbed, separately, using a window size of 100. Each data point corresponds to a masked codon. c, Two masked codons are chosen to highlight symmetric (top) and asymmetric (bottom) responses in codon preference to asymmetric perturbation. Data pertain to CaLM and synonymous perturbations.

**Figure S17:**
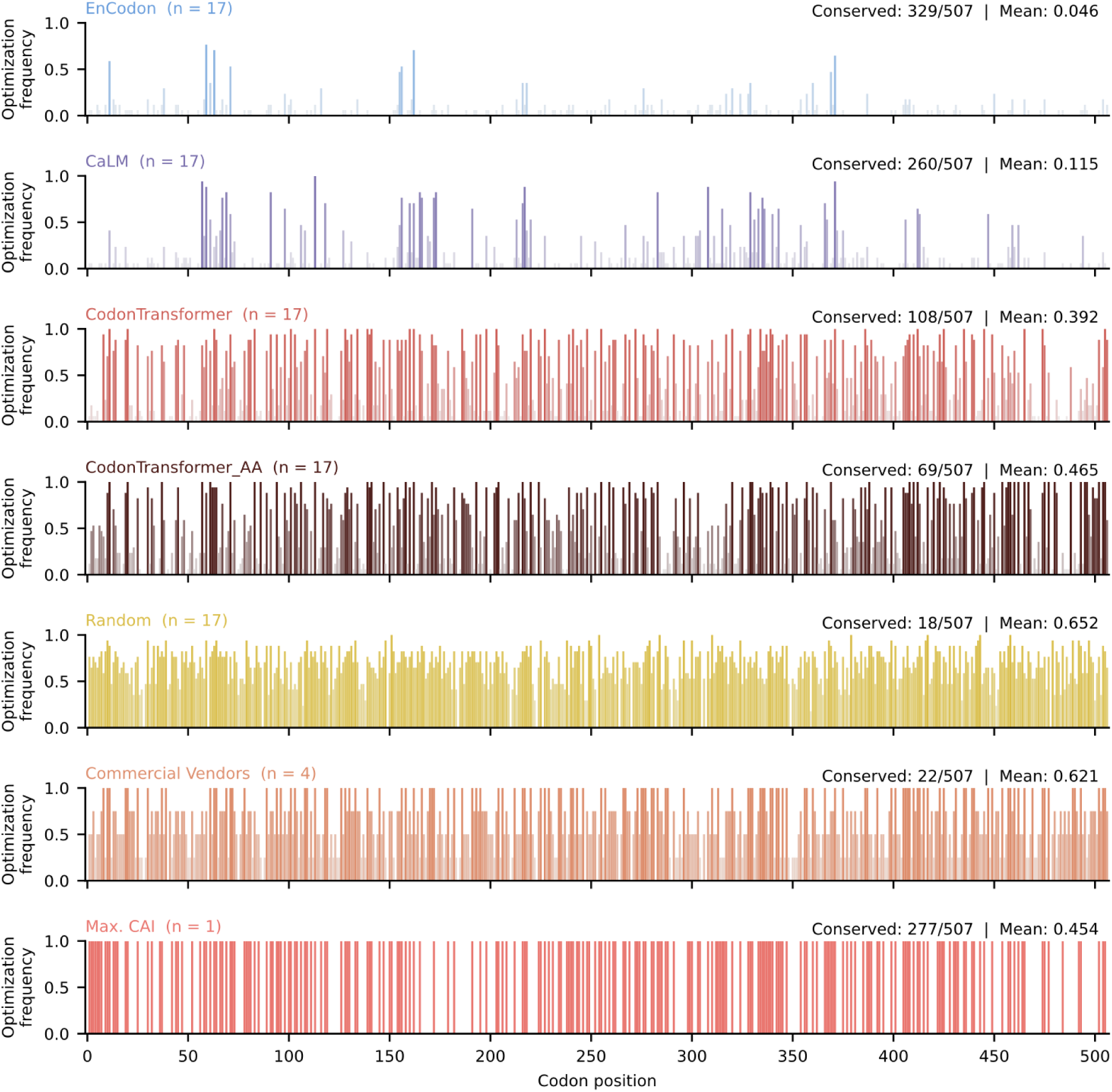
Per-position codon optimization frequency across the SEAP coding sequence by method. Each panel shows the fraction of variants within a given optimization method that differ from the reference codon at each of the 507 codon positions of the SEAP sequence. Bar intensity scales with frequency (white = fully conserved, saturated = edited in all variants). a, EnCodon retains the most reference codons (329/507 conserved, mean editing frequency 0.046), with edits concentrated at a small number of positions. b, CaLM shows a similarly conservative profile (260/507 conserved, mean editing frequency 0.115), but with more dispersed low-frequency edits. c, CodonTransformer edits more broadly (108/507 conserved, mean editing frequency 0.392), with 191 positions altered in more than half of variants. d, CodonTransformer_AA exhibits the most aggressive MLM-based recoding (69/507 conserved, mean editing frequency 0.465). e, Random synonymous substitution produces near-uniform editing across the gene (18/507 conserved, mean editing frequency 0.652), as expected from unbiased codon reassignment. f, Commercial vendor tools show a comparable editing extent to random (22/507 conserved, mean editing frequency 0.621), but with a distinct positional pattern reflecting algorithm-specific preferences. g, max. CAI (n = 1) edits 230 positions to the highest-frequency human codon, leaving 277 positions unchanged where the reference already encodes the optimal codon. Summary statistics (number of conserved positions, mean editing frequency) are annotated in each panel.

**Figure S18:**
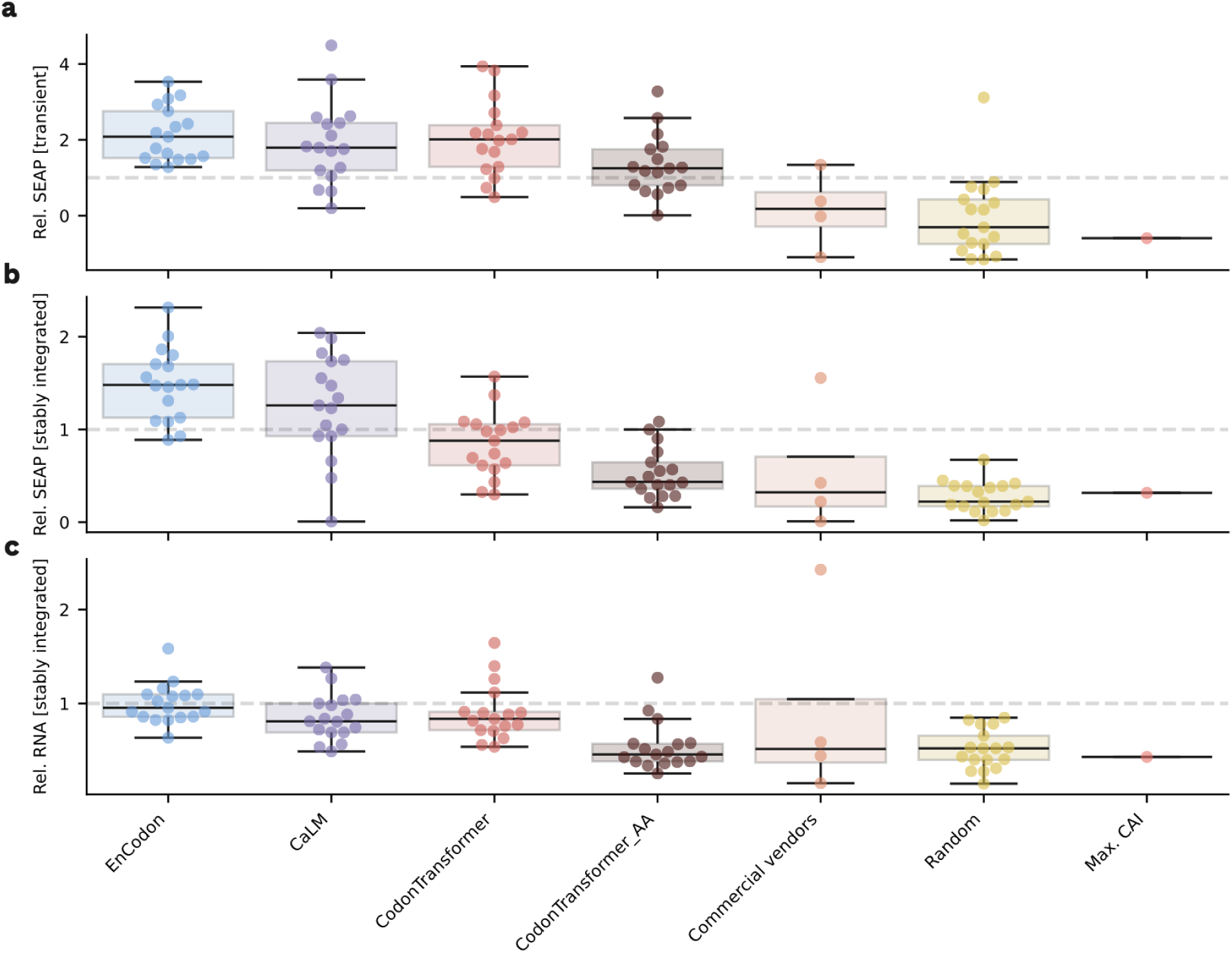
SEAP protein and mRNA expression profile across transient and stably integrated platforms. a, Relative SEAP activity following transient transfection. b, Relative SEAP activity in stably integrated cell lines. c, Relative SEAP mRNA abundance measured in stably integrated cell lines. Data are normalized relative to the Reference, indicated by the dashed grey horizontal line at y = 1.0. Individual points represent the mean of n = 3 biological replicates per sequence variant generated by the respective method (with the exception of sequence SEAP_71 rel. mRNA within the Commercial vendors group). Box plots display the mean of each respective method group across its generated sequences. The total number of data points in each plot is 92.

**Figure S19.**
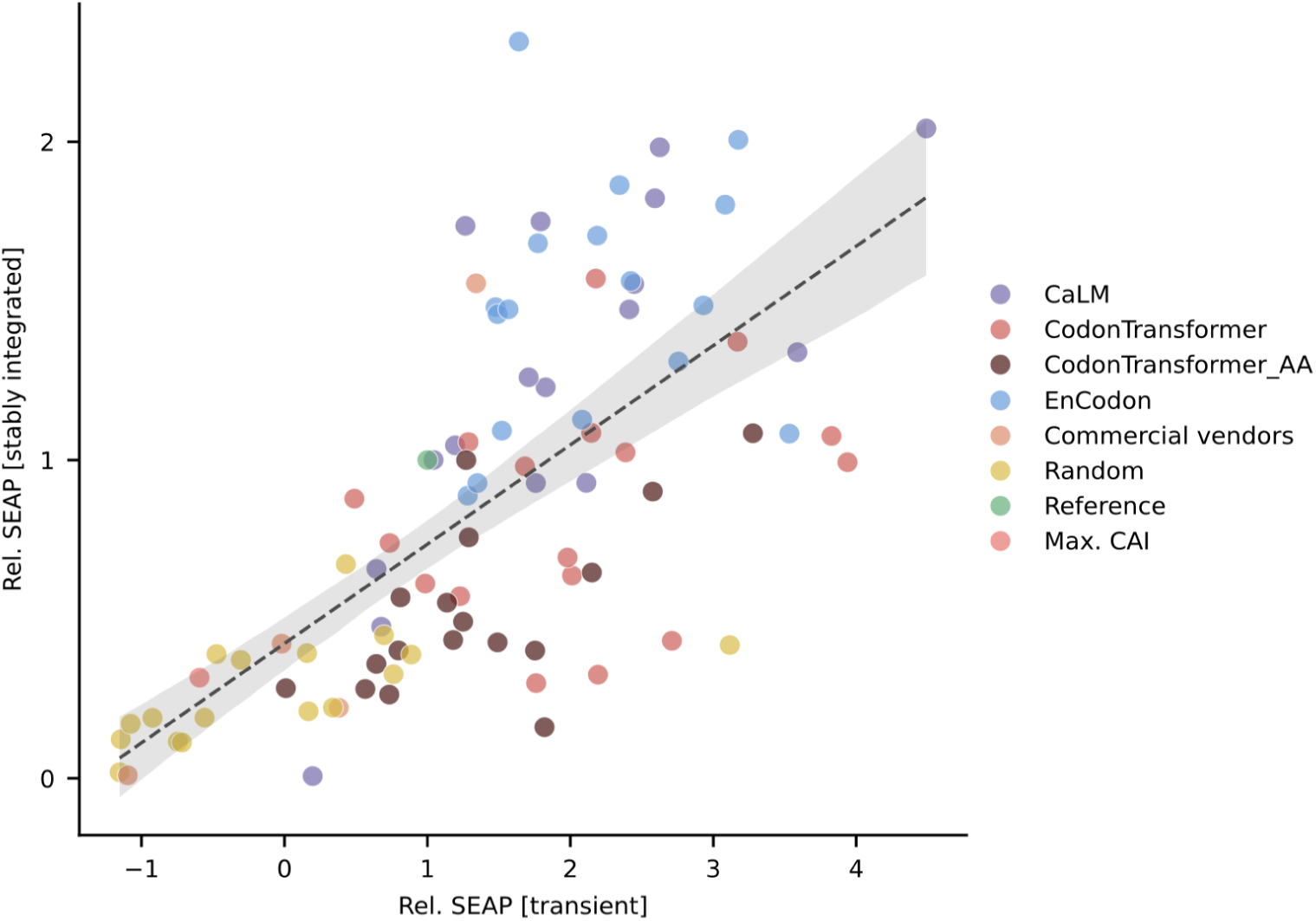
Correlation between transient and stably integrated SEAP expression. Scatter plot of normalized transient versus stably integrated SEAP activity for all 93 variants, colored by optimization method. The dashed line indicates the fit by linear regression; the shaded band represents the 95% confidence interval. Transient and stably integrated expressions are positively correlated (Spearman ρ = 0.741; R² = 0.482), indicating that relative performance differences between variants are largely maintained after genomic integration, though with attenuated magnitude. EnCodon and CaLM variants cluster in the upper right quadrant (high expression in both contexts), while random and negative control variants occupy the lower left. n = 93 variants with three distinct biological replicates.

**Figure S20.**
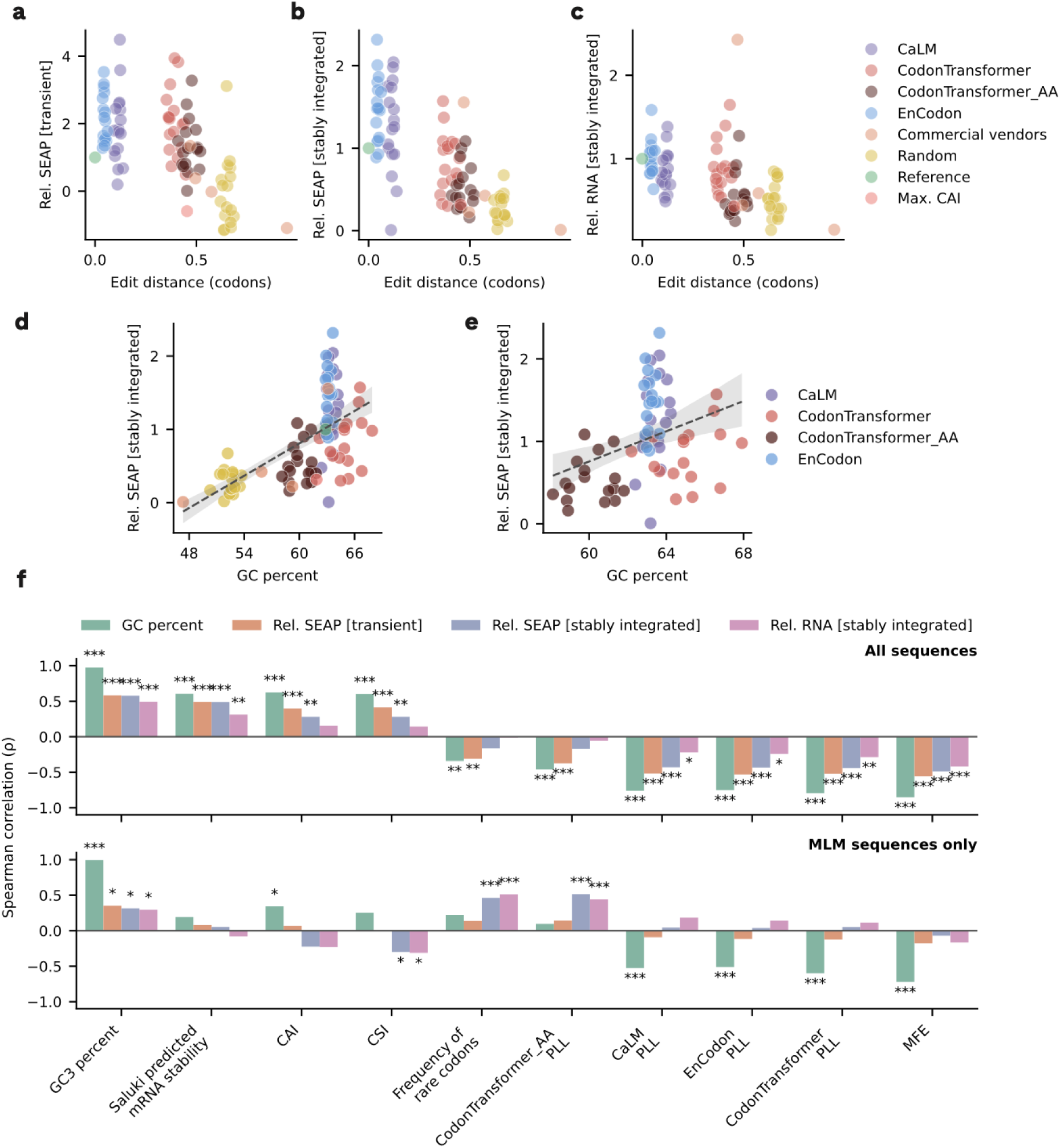
Associations between sequence features and experimental readouts. a–c, Relationship between codon-level edit distance and stably integrated SEAP expression, transient SEAP expression and RNA abundance, respectively (n=91 in each plot). d,e, Relationship between GC content and stably integrated SEAP expression across all sequence groups (n=91) (d) and within MLM-derived sequences only (n=68) (e). Dashed lines indicate fitted linear regression models, and shaded bands show 95% confidence intervals. f, Spearman correlation coefficients between sequence features and the three experimental readouts, together with GC content. Correlations were calculated across all sequence groups (top) and within MLM-derived sequences only (bottom).

**Figure S21:**
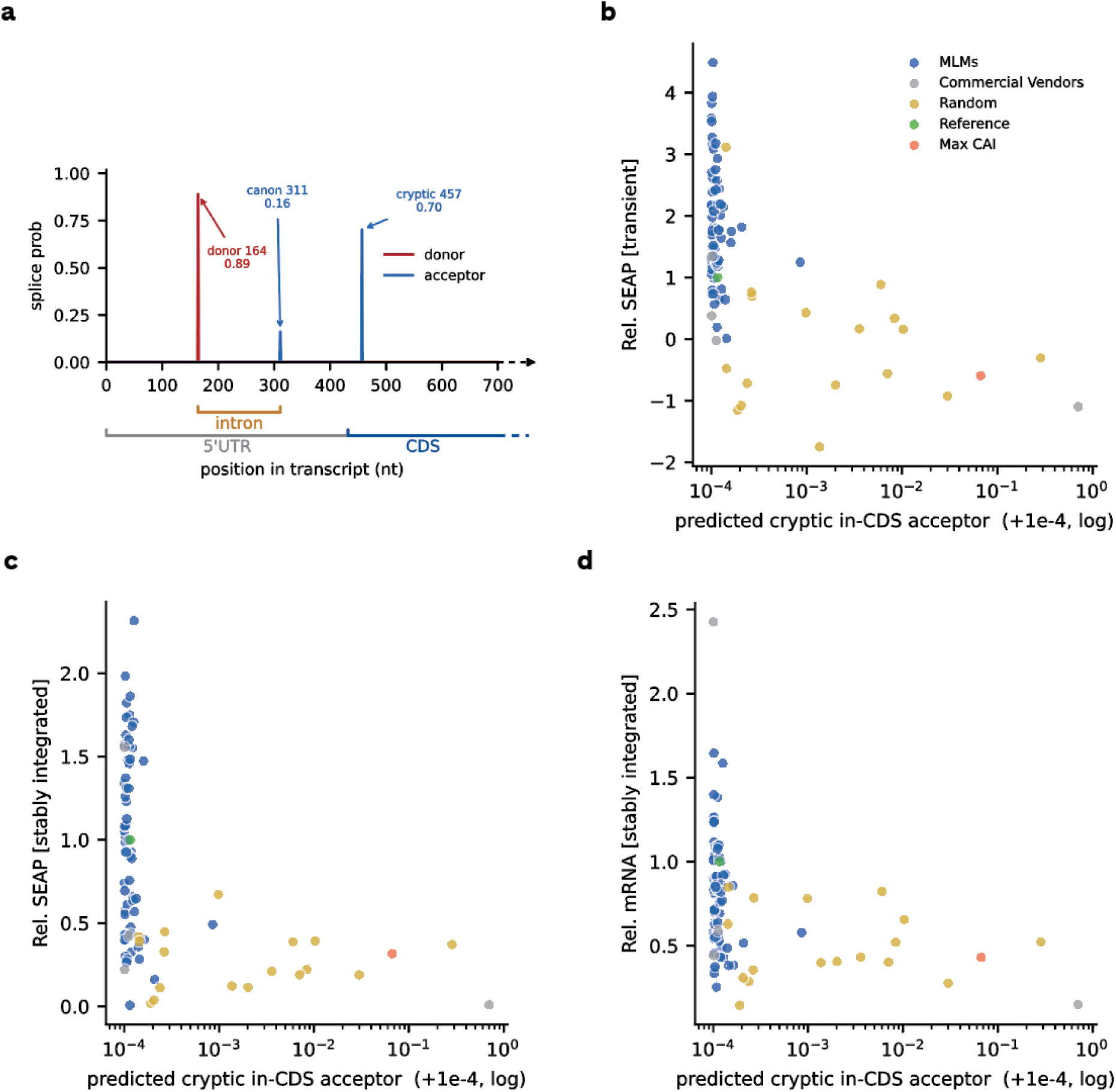
Detection of cryptic splice sites in SEAP constructs. a, Splicing donor and acceptor predictions for the (SEQID) construct obtained with OpenSpliceAI. The brackets below the x-axis indicate the boundaries between the 5’ UTR and CDS, and the position of the intron in the 5’ UTR. The site at position 457 is a cryptic splice site inside the CDS. b, relative SEAP activity following transient transfection against the maximum predicted cryptic acceptor probability in the CDS. The probability is transformed with a log(x + ε) transform, where ε = 1e-4. c, relative SEAP activity in after integration against the maximum predicted cryptic acceptor probability in the CDS. d, relative mRNA expression after integration against the maximum predicted cryptic acceptor probability in the CDS.

